# Alternative organelle targeting of OPA1 mediates fatty acid release from lipid droplets

**DOI:** 10.64898/2026.05.07.723579

**Authors:** Xiao Li, MaryEllen R. Haas, Dennis Voronin, Rikshita Bhattacharyya, Jonathon Klein, Woo Jung Cho, Camenzind G. Robinson, Robert E. Throm, Gang Wu, Chang Li, Yadav Sapkota, Natalie M. Niemi, Shondra M. Pruett-Miller, Joseph T. Opferman, Chi-Lun Chang

## Abstract

Mitochondria and lipid droplets (LDs) are functionally coupled to coordinate fatty acid utilization and storage. However, a comprehensive understanding of mitochondria–LD alliances remains elusive. We have identified a previously unrecognized role for optic atrophy 1 (OPA1), a mitochondrial fusion factor, in the regulation of fatty acid release from LDs. We demonstrated that OPA1’s exon 4 adapts an amphipathic helix to target OPA1 to LDs. OPA1 localized to LDs promotes fatty acid release by facilitating the recruitment of lipases to LDs. In addition, OPA1’s residence on LDs competes with its mitochondrial entry, influencing mitochondria fusion and connectivity. Furthermore, the S158N polymorphism within OPA1’s exon 4 exhibiting attenuated fatty acid release from LDs is associated with altered circulating lipid levels in pediatric cancer survivors. Altogether, our findings reveal that OPA1 actively mediates fatty acid release from LDs and provide a mechanistic link between OPA1 and human metabolism.

## Main

Mitochondria are the central hub for energy homeostasis, integrating products from the glycolysis-pyruvate pathway and fatty acid oxidation (FAO) into the TCA cycle to generate ATP and support energetic adaptations. Unlike pyruvate, which is readily available in the cytoplasm, fatty acids (FAs) are stored within lipid droplets (LDs) as triacylglycerols (TAGs) and cholesterol-esters^1, 2^, and must be released from the LDs prior to mitochondrial import for FAO and ATP production. Lipolysis mediates the release of FAs from LDs by coordinating lipase activity^3, 4^.

TAGs, for example, are hydrolyzed sequentially by adipose triglyceride lipase (ATGL), hormone-sensitive lipase (HSL), and monoacylglycerol lipase on the surface of LDs to become free FAs and glycerol. Lipase recruitment to LDs is central to lipolysis and regulated by protein kinase A (PKA) during active lipolytic signaling^5–8^. ATGL and HSL also play essential roles in TAG cycling, where FAs released through lipolysis are re-esterified into TAGs; this cycling is likely involved in buffering nutrient fluctuations, preventing lipotoxicity, lipid diversification, and thermogenesis^9–12^. While lipolysis and TAG cycling are well characterized in adipose tissue, increasing evidence suggests these pathways also exist across diverse cell types^13–15^. However, the molecular mechanisms governing lipase recruitment to and activation on LDs in non-adipose cell types remain poorly defined.

Cytosolic coenzyme A (CoA)-conjugated FAs can be diverted from storage and imported into mitochondria, leading to a net loss of FAs within LDs. Emerging evidence suggests that mitochondrial morphology and connectivity impact how mitochondria uptake and oxidize FAs.

Genetic perturbation promoting mitochondrial fragmentation in fibroblasts resulted in LD accumulation, a phenotype thought to arise indirectly from compromised mitochondrial FA uptake and FAO^16, 17^. Similarly, induced mitochondria elongation in mouse adipocytes was associated with enhanced FAO and reduced TAG storage^18, 19^. However, in hepatocytes, pancreatic islet β-cells, and certain lymphomas, mitochondria that have been fragmented by nutrient or genetic perturbations were found to favor FAO *in vitro*^20^. In metastatic breast cancer cells, increased FAO was accompanied by enhanced mitochondrial fission^21^. Additionally, LD-associated mitochondria with elongated morphology isolated from murine brown adipose tissues had a reduced capacity for FAO and favored TAG synthesis, leading to the expansion of LDs^22^. These apparently contradictory findings highlight a key unresolved question: how does mitochondrial morphology mechanistically influence dynamic FA storage within LDs? Notably, most existing studies have examined LD abundance only as a secondary outcome of altered mitochondrial dynamics without directly interrogating the capability of LD biogenesis and breakdown. Even though the functional coordination between LDs and mitochondria is crucial for sustaining cellular homeostasis, it is not well understood how mitochondria coordinate FA release from LDs.

One such ‘missing’ coordinator may be optic atrophy 1 (OPA1); OPA1 is a well characterized and broadly expressed mitochondrial morphological factor that promotes inner mitochondrial membrane fusion via its GTPase activity^23^. In humans, eight OPA1 mRNA isoforms are produced by alternative splicing of exons 4, 4b, and 5b^24^. Following translation and mitochondrial import, OPA1 is anchored on the inner membrane as the long OPA1 (L-OPA1) isoforms, which are subjected to further proteolytic processing to generate soluble, short OPA1 (S-OPA1) isoforms in the intermembrane space^25^. Whether inner membrane fusion can be mediated by L-OPA1 alone, or instead requires both L-OPA1 and S-OPA1, remains an open question^26–30^. In addition to its roles within mitochondria, some evidence suggests OPA1 may have non-canonical functions regulating lipid storage. For example, OPA1 has been detected on LDs in cultured adipocytes and adrenal gland cells; however, whether it plays a role in PKA-mediated signaling appears to be cell-type dependent^31, 32^. Multiple proteomic studies have also identified OPA1 in the LD proteome in mouse skeletal muscle, cultured hepatocytes, and U2OS and HEK293 cell lines^33–35^. In mice, OPA1 promotes white adipose tissue expansion and browning^36, 37^ and is required for lipid metabolism and cold-induced thermogenic activation^37, 38^. Despite these findings, the precise molecular mechanisms by which OPA1 orchestrates LD remodeling, and whether these mechanisms extend to non-adipose cells, are unknown.

Here, we report that OPA1 mediates FA release via exon 4-directed LD targeting. Through comprehensive analyses of FA fluxes, we demonstrate that, in addition to its canonical functions in mitochondria, OPA1 actively regulates FA release from LDs via two interrelated mechanisms: (1) first, exon 4 adopts an amphipathic helix that directs a subpopulation of OPA1 to LDs. A naturally occurring OPA1 isoform 2, which lacks exon 4, is exclusively distributed to mitochondria. (2) Second, LD-associated OPA1 facilitates the recruitment of ATGL and HSL to LDs under resting conditions. The recruitment of these lipases and the consequent FA release positively correlate with the level of OPA1 observed on LDs. When residing on LDs, OPA1’s mitochondrial localization is effectively reduced, ultimately abating mitochondria fusion and connectivity. Notably, a common human polymorphism (*i.e.,* S158N within exon 4) decreases OPA1 targeting to LDs and is associated with altered circulating lipid levels from a database of cancer survivors. Altogether, our findings reveal a new role for the ubiquitously expressed OPA1 – mediating FA release from LDs – and suggest a mechanistic link between LD metabolism and mitochondrial morphology through OPA1’s alternative organelle targeting.

## Results

### Defective FA release from LDs in OPA1 knockout cells

FA metabolism is regulated by dynamic storage and release at LDs and mitochondrial FA consumption (Fig. 1A). To investigate how OPA1 impacts LDs, we used a battery of assays with pharmacological inhibitors to independently evaluate FA release from LDs, mitochondrial FA import, and mitochondrial FAO in wildtype (WT) and *OPA1* knockout (KO) U2OS cells generated via CRISPR-genome editing and validated by sequencing, western blotting, and immunostaining (Fig. S1A–C). We first established a FA release assay by subjecting cells to oleic acid (OA) withdrawal in the presence of triacsin C (Fig. 1B, top panel), an inhibitor of long FA acyl-CoA synthases, to blunt acyl-CoA–dependent LD biogenesis and acylcarnitine synthesis required for mitochondrial FA import simultaneously^39–42^. By eliminating LD biogenesis and mitochondrial FA consumption, the changes in existing LD content can be attributed primarily to FA release from LDs, allowing us to assess the net FA release. In U2OS cells that were first treated with 100 µM OA overnight to induce LD accumulation before incubation with triacsin C in the absence of OA (withdrawal) for 6 h, we observed a significant reduction (21% remaining) in LD content in WT cells (Fig. 1B,C). We confirmed the triacsin C-mediated reduction was indeed FA release from LDs as co-incubation with triacsin C and NG497, an ATGL inhibitor^43^, but not co-incubation with triacsin C and Lalistat 2, a lysosomal lipase inhibitor^44^, prevented the loss of LDs (Fig. S2A). Co-incubation of triacsin C and H89, a PKA inhibitor^45^, minimally affected LD breakdown (Fig. S2B), suggesting that PKA did not participate in the triacsin C–mediated FA release in resting state U2OS cells. Intriguingly, when comparing *OPA1* KO and WT U2OS cells with similar baseline LD content following OA incubation overnight, the *OPA1* KO cells displayed significantly higher LD accumulation (51% remaining) than WT cells following triacsin C treatment (Fig. 1B,C), suggesting defects in FA release from LDs. This defective FA release was also observed in *OPA1* knockdown HeLa cells and *OPA1* KO mouse embryonic fibroblasts (MEFs) (Fig. S2C–F), indicating OPA1’s role in regulating FA release may be generalizable across different cell types. Consistent with these observations, we found *OPA1* KO U2OS cells retained significantly higher quantities of LDs than WT cells following glucose starvation for 20 h (60% in KO vs 34% in WT; Fig. S2G). To continue this characterization, we introduced yellow fluorescent protein (YFP)-tagged OPA1 isoform 1 (OPA1-YFP), the most abundant OPA1 isoform, into the *OPA1* KO cells via lentiviral transduction. We found that the accumulated LD content in *OPA1* KO cells after the 6-h triacsin C incubation, was significantly reduced by OPA1-YFP expression (Fig. 1D).

**Figure 1.**
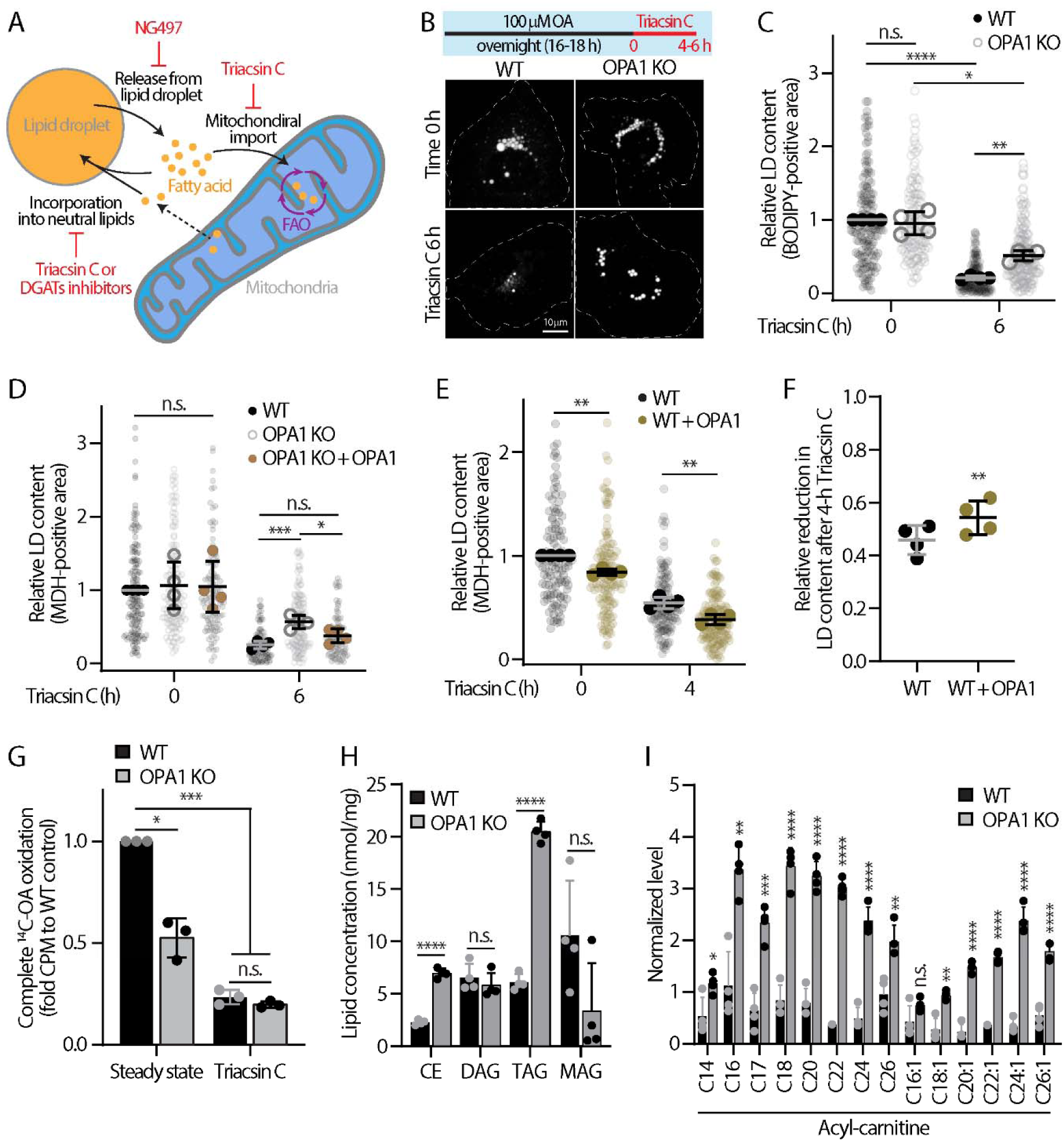
Comprehensive analysis of fatty acid trafficking reveals defective release from lipid droplets in *OPA1* knockout cells. **(A)** Multidirectional fatty acid (FA) trafficking pathways between lipid droplets (LDs) and mitochondria. Terms: FAO, fatty acid oxidation; NG497, an ATGL inhibitor; triacsin C, an inhibitor of long FA acyl-CoA synthetase. **(B)** A timeline illustrating oleic acid (OA) withdrawal experiment (top) and confocal images (bottom) of BODIPY 493/503-labeled LDs before and after 6-h incubation with 10 µM triacsin C in wildtype (WT) and *OPA1* knockout (KO) U2OS cells treated with 100 µM OA overnight. Maximal intensity projected confocal images from whole cells are shown. Dashed lines mark cell boundary. **(C)** Quantification of relative BODIPY 493/503-positive LD content as described in (B). Mean ± standard deviation from four independent experiments are shown (n = 148–200 cells). n.s., no significance, ****P ≤ 0.0001, **P ≤ 0.01, *P ≤ 0.05, as assessed by one-way ANOVA. **(D)** Quantification of relative MDH-positive LD content in WT, *OPA1* KO, and OPA1-expressing *OPA1* KO U2OS cells treated with 100 µM OA overnight followed by 10 µM triacsin C incubation for 6 h. Mean ± standard deviation from four independent experiments are shown (96–148 cells). n.s., no significance, ***P ≤ 0.001, *P ≤ 0.05, as assessed by one-way ANOVA. **(E)** Quantification of MDH-positive LD content in WT and OPA1-expressing WT U2OS cells treated with 100 µM OA overnight followed by 10 µM triacsin C incubation for 4 h. Mean ± standard deviation from four independent experiments are shown (124–141 cells). n.s., no significance, **P ≤ 0.01, as assessed by one-way ANOVA. **(F)** Quantification of the reduction in relative LD content after 4-h triacsin C treatment as shown in (E). Mean ± standard deviation from four independent experiments are shown. **P ≤ 0.01, as assessed by paired *t*-test. **(G)** Scintillation counts per minute (CPM) for complete ^14^C-OA oxidation in WT and *OPA1* KO U2OS cells under steady state and Triacsin C-treated conditions. Mean ± standard deviation from three independent experiments are shown. n.s., no significance, ***P ≤ 0.001, *P ≤ 0.05, as assessed by one-way ANOVA. **(H)** Concentration of cholesterol-ester (CE), diacylglycerol (DAG), triglyceride (TAG), and monoacylglycerol (MAG) in steady-state WT and *OPA1* KO U2OS cells determined via liquid chromatography–mass spectrometry. Mean ± standard deviation from four replicates are shown. n.s., no significance, ****P ≤ 0.0001, as assessed by unpaired *t*-test. **(I)** Variance in acyl-carnitine levels across acyl-chain length in steady-state WT and *OPA1* KO U2OS cells determined using liquid chromatography–mass spectrometry. Mean ± standard deviation from four replicates are shown. n.s., no significance, ****P ≤ 0.0001, ***P ≤ 0.001, **P ≤ 0.01, *P ≤ 0.05, as assessed by unpaired *t*-test.

Additionally, WT cells overexpressing OPA1 exhibited less LD content than control WT cells under both OA-fed and 4-h triacsin C–treated conditions (Fig. 1E), with a more pronounced reduction in LD content following triacsin C incubation (Fig. 1F), suggesting that OPA1 overexpression promotes LD consumption. These results suggest OPA1 plays a direct role mediating FA release and the observed changes are not secondary effects of OPA1’s KO. Furthermore, we found comparable levels of expression in key enzymes (*e.g.,* ATGL, HSL, and CGI-58, which is a co-activator for ATGL^46–48^) involved in LD breakdown in WT and *OPA1* KO cells (Fig. S1D,E), suggesting that the observed FA release defect in *OPA1* KO cells was not due to compensatory mechanisms associated with aberrant expressions of these key enzymes.

We continued to validate that the *OPA1* KO U2OS cells we generated faithfully recapitulated canonical fatty acid–related phenotypes by first functionally validating FAO capability using ^14^C-labeled OA. We found FAO was significantly lower at steady state in *OPA1* KO cells than in WT cells, while triacsin C treatment, used as a negative control, caused a significant reduction in FAO in both populations (Fig. 1G). These findings are consistent with previous studies demonstrating OPA1 loss impaired FAO^49, 50^.

To determine if mitochondrial FA import attributed the defective FAO we observed, we transiently pulsed cells with 1 µM fluorescently labeled FAs for 10 min in the presence of DGAT1 and DGAT2 inhibitors to eliminate LD biogenesis, which would compete with mitochondria for FAs. Following the transient pulse, TopFluor (TF)-OA signal was readily detected in mitochondria in both WT and *OPA1* KO U2OS cells, regardless of their mitochondrial morphology (Fig. S2H). Quantification of TF-OA intensity in mitochondria showed 20% less FA import in *OPA1* KO cells compared to WT cells (Fig. S2I), likely contributing to the observed defects in FAO. Triacsin C treatment, used as a negative control, blunted TF-OA uptake in both WT and *OPA1* KO cells (Fig. S2I). Coincidently, import of BODIPY-linoleic acid (LA) was similar while import of NBD-arachidonic acid (AA) was moderately enhanced in *OPA1* KO cells (Fig S2I), suggesting differential regulation of FA import. Altogether, loss of OPA1 severely limited FAO while impacting FA entry into mitochondria differently depending on the type of FA, underlining that FAO and mitochondrial FA import can be independently regulated.

To corroborate the observed defective FA release from LDs and mitochondrial FAO, we performed lipidomic and metabolomic analyses in WT and *OPA1* KO U2OS cells in steady state. TAG and cholesterol-ester levels were ∼3-fold higher in *OPA1* KO cells than in WT cells (Fig. 1H). *OPA1* KO cells also displayed moderately, but not significantly, lower monoacylglycerol levels, a broken-down product from TAG hydrolysis by lipases, than the WT (Fig. 1H). Additionally, overall lipid synthesis was not strongly affected in the *OPA1* KO cells, though several phospholipid species were moderately upregulated (Fig. S2J,K). Furthermore, nearly all detected acyl-carnitine species levels were significantly higher in *OPA1* KO cells than in WT cells (Fig. 1I), reflecting defective FAO in the *OPA1*-depleted mitochondria. Altogether, our data demonstrate that OPA1, in addition to its canonical functions in mitochondrial morphology and FAO, can independently impact FA release from neutral lipids within LDs in the absence of lipolytic signaling.

### A subpopulation of OPA1 localizes to LDs

To investigate how mitochondrial OPA1 might regulate FA release from LDs, we compared OPA1-YFP localization in control and OA-treated (LD-laden) WT U2OS cells. Using confocal microscopy, we found OPA1 exclusively colocalized with mitochondria in untreated cells, while a subpopulation of OPA1-YFP resided on LDs in OA-treated cells (Fig. 2A). Airyscan microscopy with enhanced spatial resolution^51^ and line scanning analysis of OA-treated cells further revealed that OPA1-YFP decorated LDs without mitochondrial association (Fig. S3A,B), and this LD-associated OPA1-YFP was highly colocalized with a LD-resident protein, perilipin 2 (PLIN2) (Fig. S3C,D). We quantified the relative enrichment of OPA1 on LDs and found a significant increase in LD enrichment in OA-treated WT U2OS cells; as well as in cells incubated with LA or AA to induce LD formation (Figs. 2B and S3E), suggesting that the FA content within LDs is not a crucial determinant for OPA1’s localization. Consistent with this observation, a subpopulation of OPA1-YFP was also found on LDs in HeLa and Huh7 cells treated with OA overnight (Fig. S3F). In addition, a small population of endogenous OPA1 was observed on clusters of small LDs in U2OS cells (Fig. 2C,D) using an OPA1 antibody validated in *OPA1* KO U2OS cells (Fig. S1C). Subcellular fractionation further revealed that endogenous OPA1, especially L-OPA1 (top high-molecular weight bands), was enriched in the LD fraction as validated by the LD protein PLIN2, while S-OPA1 (lower bands) generated via mitochondrial proteolysis was primarily present in the cytosol and mitochondria-enriched pellet fractions (Fig. 2E). Prohibitin 1 (PHB1), a mitochondrial inner membrane protein, was chosen as our mitochondrial marker because it localizes to the intermembrane space^52^, the same compartments that OPA1 distributes to canonically.

**Figure 2.**
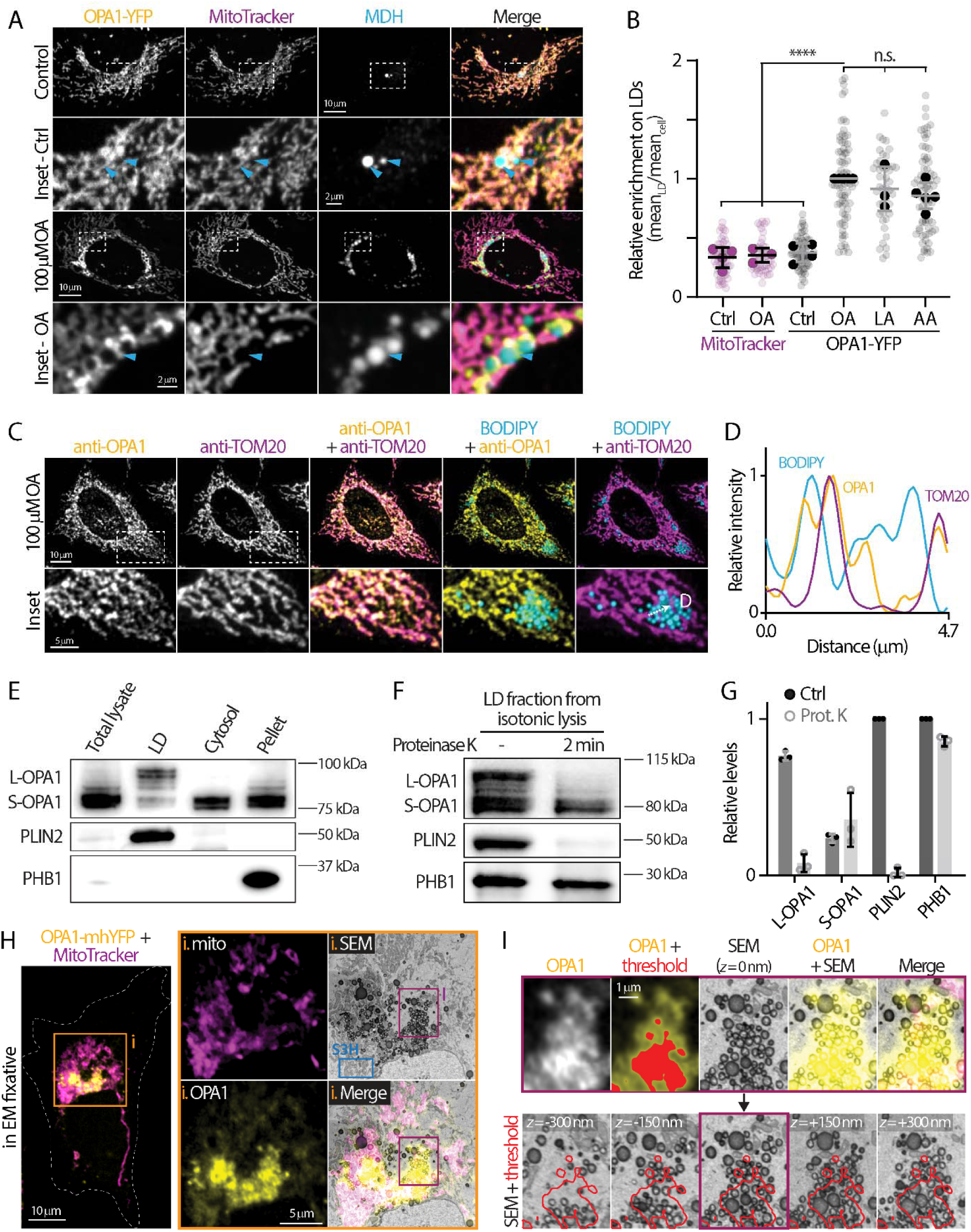
A subpopulation of OPA1 is localized to lipid droplets. **(A)** Localization of OPA1-YFP (lentiviral), mitochondria (MitoTracker Deep Red), and lipid droplets (LDs; MDH) in U2OS cells ± overnight 100 µM oleic acid (OA) treatment. Maximal intensity projected confocal images from four axial slices (∼1 µm total thickness) are shown. Cyan arrowheads indicate area with LDs. **(B)** Relative enrichment of MitoTracker and OPA1 on LDs from (A) and in cells treated with 100 µM linoleic acid (LA) or arachidonic acid (AA) overnight. Means ± standard deviation from 3–5 independent experiments are shown (total of 43–74 cells). n.s., no significance, ****P ≤ 0.0001, as assessed by one-way ANOVA. **(C)** Subcellular localization of endogenous OPA1 on mitochondria (anti-TOM20) and LDs labeled by BODIPY 493/503 in an OA-treated U2OS cell monitored with confocal microscopy. Sum of confocal images from five axial slices (∼1.2 µm in total thickness) are shown. **(D)** Relative intensity profiles of OPA1, TOM20, and BODIPY measured along the white-dashed arrow from lower left panel in (C). **(E)** Western blot analysis of endogenous OPA1, PLIN2 (a LD protein), and PHB1 (a mitochondrial inner membrane protein) in sucrose-gradient cellular fractionations from U2OS cells treated with 500 μM OA. **(F)** Western blot analysis of OPA1, PLIN2, and PHB1 in isolated LD fractions from U2OS cells before and after treatment with 10 µg/ml of proteinase K treatment for 2 min. **(G)** Quantification of OPA1, PLIN2, and PHB1 levels in the proteinase K protection assay as shown in (F). Means ± standard deviation from three independent experiments are shown. **(H)** Correlative confocal-scanning electron microscopy (SEM) images of OPA1 on LDs and in mitochondria in OPA1-mhYFP–expressing U2OS cells treated with 100 µM OA overnight. Confocal and SEM images from a single axial slice are shown. EM, electron microscopy. Insets outlined in purple and blue are shown in (I) and Fig. S3H, respectively. **(I)** Five serial SEM images along the z-axis at 150-nm step are shown. The dark red box marks the region with higher OPA1-YFP fluorescence signal as the threshold.

To address the possibility that the L-OPA1 enrichment observed in the LD fraction (Fig. 2E) originated from a small number of mitochondria co-purified with LDs, we performed a proteinase K protection assay using LD fractions purified via isotonic lysis to ensure mitochondrial membrane integrity. This assay allowed us to distinguish between OPA1 resulting from mitochondrial contamination (protected from digestion by the mitochondrial outer membrane) and OPA1 localized to LDs (subjected to digestion). We first confirmed the presence of OPA1, PLIN2, and PHB1 in the isolated LD fraction, indicating a low degree of co-purification of mitochondria and LDs. Following proteinase K incubation, the amount of L-OPA1, but not S-OPA1 was greatly reduced. Similarly, LD-resident protein PLIN2 was completely degraded, while PHB1, which localizes to the intermembrane space, was protected from digestion (Fig. 2F,G). L-OPA1 in LD fractions isolated from mouse livers were also subjected to proteinase K digestion while PHB1 remained protected (Fig. S3G). These observations demonstrated that, despite potential mitochondrial contamination in isolated LD fractions, the majority of L-OPA1 is most likely associated with LDs directly, or it would not have been susceptible to digestion in the proteinase K protection assay (Fig. 2F,G).

We next performed correlative confocal-scanning electron microscopy (SEM) to validate these observations with high spatial resolution. We expressed a monomeric hyperfolder-YFP tagged OPA1 (OPA1-mhYFP) that retains fluorescence in electron microscopy fixatives^53^ in U2OS cells stained with MitoTracker as a fiducial marker for correlative SEM. OPA1-mhYFP signals were found in regions enriched with mitochondria (Fig. S3H) and in areas containing small LDs (Fig. 2H; orange box). We focused on the strongest OPA1 fluorescence signal (Fig. 2I; threshold in red) and reviewed the corresponding region in the correlative serial SEM images. Using a 150-nm step size throughout the entire 600-nm thickness, which is similar to confocal axial resolution, we found the corresponding regions to be composed of LD clusters with no detectable mitochondria (Fig. 2I, bottom panels), suggesting that a subpopulation of OPA1 resides on the surface of LDs.

### An alternative LD-targeting motif competes with OPA1 mitochondrial import

OPA1 is a nuclear-encoded protein that was not detected in previous co-translationally inserted mitochondrial proteomes^54, 55^, suggesting it may be translated in the cytosol prior to reaching its organelle destination. To interrogate the timing of OPA1’s LD targeting following its translation, we introduced an inducible OPA1-YFP (iOPA1-YFP) under a tetracycline-driven promoter into *OPA1* KO U2OS cells to avoid endogenous OPA1-directed distribution via dimerization. The iOPA1-YFP signals were detected on both mitochondria and LDs after ∼4 h of doxycycline induction (Fig. 3A). Although the signal was dim, we were able to categorize the iOPA1-YFP signal based on its first detected organelle distribution and found that on average 45% of the cell population had iOPA1 exclusively in mitochondria, 33% had it exclusively on LDs, and 22% showed the signal in both organelles (Fig. 3B).

**Figure 3.**
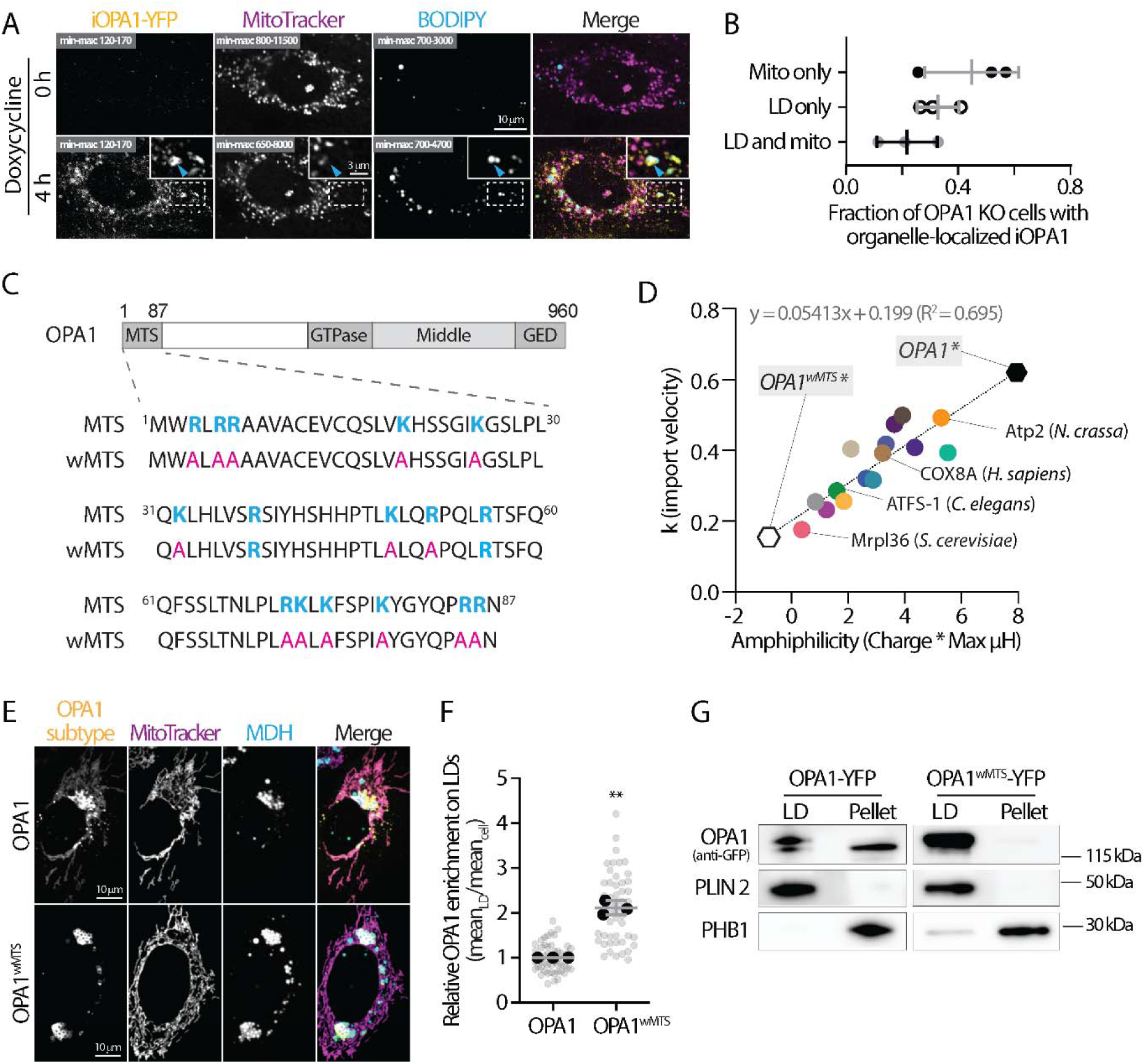
An alternative lipid droplet–targeting motif competes with OPA1 mitochondrial import. **(A)** Subcellular distribution of inducible OPA1 (iOPA1)-YFP on lipid droplets (LDs; BODIPY 665/676) and in mitochondria (MitoTracker Red) in *OPA1* knockout (KO) U2OS cells following treatment with 4 µg/mL doxycycline for 4 h. Confocal images from a single axial slice are shown. Cyan arrowheads indicate regions containing LDs. **(B)** Fraction of cells with iOPA1 localized to indicated organelle as described in (A). Means ± standard deviation from three independent experiments are shown. **(C)** Schematic illustrating the mitochondrial targeting sequences (MTSs) of OPA1 and OPA1 mutations that result in a weakened MTS (wMTS). Positively charged amino acid residues are indicated with bold blue; alanine substitutions are shown in pink. Amino acid number and protein domains are indicated. Abbreviation: GED, GTPase effector domain. **(D)** Correlation between the calculated amphiphilicity and experimentally determined import velocity (k) of MTSs from 16 mitochondrial proteins (listed in Fig. S4A). Import velocity of OPA1’s MTS and its mutated wMTS were predicted (marked by asterisk) based on the liner regression formula. Example data points of Atp2 (*Neurospora crassa*), COX8A (*Homo sapiens*), ATFS-1 (*Caenorhabditis elegans*) and Mrpl36 (*Saccharomyces cerevisiae*) are indicated. (**E**) Localization of OPA1-YFP and OPA1^wMTS^-YFP, mitochondria (MitoTracker Deep Red), and LDs (MDH) in U2OS cells treated with 100 µM OA overnight detected by confocal microscopy. Maximal intensity projected confocal images from four axial slices (∼1 µm total thickness) are shown. **(F)** Relative enrichment of OPA1 and OPA1^wMTS^ on LDs in U2OS cells treated with 100 µM oleic acid overnight in (E). Means ± standard deviation from three independent experiments are shown (n = 51 cells/condition). **P ≤ 0.01, as assessed by unpaired *t*-test. **(G)** Western blot analysis of OPA1-YFP, or OPA1^wMTS^-YFP, PLIN2 and PHB1 in isolated LD and mitochondria-enriched pellet fractions from U2OS cells.Results from one out of three independent experiments are shown.

Dual localization of mitochondrial proteins has been linked to a ‘weak’ mitochondrial targeting sequence (MTS), which disallows efficient mitochondrial import to promote alternative subcellular localization^56, 57^. To determine if OPA1 has a weak MTS facilitating its localization at LDs, we used OPA1’s previously mapped MTS (87 amino acids in length^28^, Fig. 3C) and calculated its amphiphilicity, a product of the charge and maximal hydrophobic moment of a MTS, and found it had the highest score compared with 16 experimentally determined MTSs across proteins from multiple species (Figs. 3D and S4A), suggesting a strong import behavior^58, 59^. This suggests OPA1’s LD localization is not due to a ‘weak’ MTS responsible for inefficient mitochondrial import, but may instead result from inclusion of a LD-targeting motif that competes with the MTS to direct L-OPA1 to LDs shortly following its production in the cytoplasm. To test this hypothesis, we generated an OPA1 variant with a weakened MTS OPA1^wMTS^, by substituting most of the positively charged residues within its MTS with alanine (Fig. 3C). OPA1^wMTS^ was predicted to have substantially reduced mitochondria import efficiency (Figs. 3D and S4A), and consistent to this prediction, displayed a pronounced cytosolic distribution in addition to mitochondria localization in U2OS cells under resting condition (Fig.S4B, Cell 1). Cells with a subpopulation of OPA1^wMTS^ decorating LDs were also found in resting state cells (Fig. S4B, Cell 2), resulting in a greater enrichment of OPA1^wMTS^ on LDs than OPA1 on LDs in the absence of OA (Fig. S4C). In OA-treated (LD-laden) U2OS cells, OPA1^wMTS^-YFP was found predominantly on LDs with minimal mitochondrial distribution (Fig. 3E,F). OPA1^wMTS^’s higher affinity for LDs was further underlined by subcellular fractionation, as we detected more OPA1^wMTS^ in isolated LD fractions than we did WT OPA1-YFP, relative to their mitochondria pellet fractions (Fig. 3G). Altogether, these results demonstrate that weakening mitochondrial OPA1 import promotes OPA1’s localization to LDs, providing support for a model in which OPA1 possesses competing targeting motifs (*i.e.,* a MTS vs a LD-targeting motif) that direct its partitioning between mitochondria and LDs.

### OPA1’s exon 4 constitutes an amphipathic helix for LD targeting

To identify OPA1’s LD-targeting motif, we generated YFP-tagged, truncated forms of OPA1 focusing on its first 195 amino acids, which are exclusively present in L-OPA1 (Fig. 4A)^28^. OPA1 with its MTS deleted (OPA1^ΔMTS^) only distributed to LDs (Fig. 4B,C), indicating the MTS is not required for LD targeting and an additional LD-targeting motif must reside within amino acids 88–195. While fragments of OPA1^88–195^ and OPA1^127–195^ primarily localized to LDs, OPA1^157–195^, with a truncated exon 4, became cytosolic (Figs. 4A and S5A). This observation prompted us to hypothesize that exon 4 is required for OPA1’s distribution to LDs.

**Figure 4.**
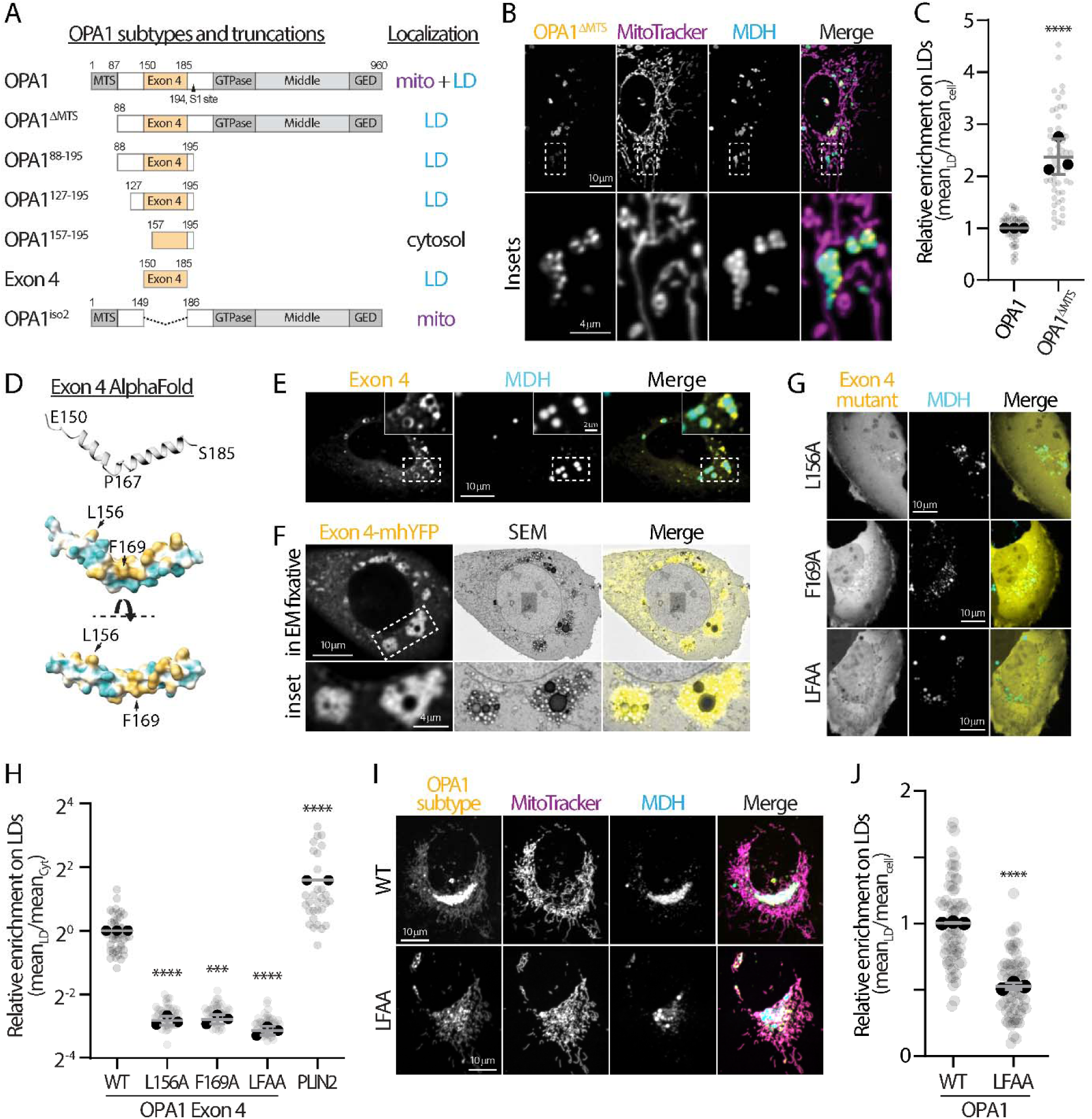
OPA1’s exon 4 adapts an amphipathic helix structure to target lipid droplets. **(A)** Constructs expressing full-length and truncated OPA1, as well as summary of their primary subcellular localizations. Amino acid number and protein domains are indicated. Abbreviations: MTS, mitochondria targeting sequence; GED, GTPase effector domain; LD, lipid droplet; mito, mitochondria. **(B)** Localization of OPA1^ΔMTS^-YFP, mitochondria (MitoTracker Deep Red), and lipid droplets (MDH) in U2OS cells treated with 100 µM oleic acid (OA) overnight and detected by confocal microscopy. Maximal intensity projected (MIP) confocal images from four axial slices (∼1 µm total thickness) are shown. **(C)** Relative enrichment of OPA1^ΔMTS^ from (B) and OPA1 on LDs in U2OS cells treated with 100 µM OA overnight. Means ± standard deviation from three independent experiments are shown (total of 48–55 cells). ****P ≤ 0.0001, as assessed by unpaired *t*-test. **(D)** Amphipathic helix structure of exon 4 as predicted by AlphaFold. Cyan and yellow indicate polar and non-polar amino acids, respectively, and black arrows indicate the location of residues for mutagenesis. **(E)** Localization of YFP-tagged exon 4 in MDH-stained U2OS cells treated with 100 µM OA overnight and detected by confocal microscopy. MIP confocal images from four axial slices (∼1 µm total thickness) are shown. **(F)** Correlative confocal-scanning electron microscopy (SEM) images of exon 4 on LDs in exon 4-mhYFP expressing U2OS cells treated with 100 µM OA overnight. Confocal and SEM images from a single axial slice are shown. **(G)** Localization of exon 4 mutants relative to LDs (MDH) in U2OS cells treated with 100 µM OA overnight and detected by confocal microscopy. MIP confocal images from four axial slices (∼1 µm total thickness) are shown. LFAA, a double mutation of L156 and F169 to alanines. **(H)** Relative enrichment of exon 4 and mutants (described in E and G), as well as PLIN2-GFP on LDs in U2OS cells treated with 100 µM OA overnight. Means ± standard deviation from two or three independent experiments are shown (total of 30–52 cells). ****P ≤ 0.0001, ***P ≤ 0.001, as assessed by one-way ANOVA. **(I)** Localization of OPA1 and OPA1^LFAA^, mitochondria (MitoTracker Deep Red), and LD (MDH) in U2OS cells treated with 100 µM OA overnight and detected by confocal microscopy. MIP confocal images from four axial slices (∼1 µm total thickness) are shown. **(J)** Relative enrichment of OPA1 and OPA1^LFAA^ from (I) on LDs in cells treated with 100 µM OA overnight. Means ± standard deviation from three independent experiments are shown (total of 63–65 cells). ****P ≤ 0.0001, as assessed by one-way ANOVA.

AlphaFold 3 predicts that exon 4 adopts a secondary structure of an amphipathic helix with proline^1^^67^ close to the center slightly bending the helix (Fig. 4D). Exon 4 is located within one of OPA1’s unstructured regions that is readily exposed to the cytoplasm (AlphaFold 3 prediction; data not shown). The non-polar residues (Fig. 4D, yellow) are slightly offset along the long axis, suggesting a weak interaction with membranes. Nonetheless, overexpressed exon 4-YFP displayed clear LD localization in WT U2OS cells (Fig. 4E). Correlative confocal-SEM imaging revealed that exon 4-mhYFP localized to clusters containing LDs of various sizes (Fig. 4F). By mutating specific non-polar amino acids (i.e., leucine^1^^56^, phenylalanine^1^^69^, or both) to alanine completely disrupted exon 4’s ability to localize to LDs (Fig. 4G,H). Consistent with this observation, full-length OPA1^LFAA^ (OPA1 with the double mutation of leucine^1^^56^ and phenylalanine^1^^69^ to alanines) also showed significantly lower enrichment on LDs than WT OPA1 (Fig. 4I,J), emphasizing OPA1’s ability to distribute to LDs likely requires these non-polar residues. Intriguingly, exon 4’s affinity for LDs was significantly lower than the LD-resident protein PLIN2’s (Fig. 4H), suggesting a dynamic interaction between LDs and OPA1 via exon 4 coordinates OPA1’s mitochondrial import and canonical functions.

Since OPA1 contains both LD- and mitochondria-targeting motifs, we considered the possibility that OPA1 bridges mitochondria and LD at contact sites. To test this hypothesis, we overexpressed OPA1 in WT U2OS cells and found the colocalization between LD and mitochondria was not affected (Fig. S5B). In addition, the levels of LD-mitochondria contact sites remained comparable in WT and *OPA1* KO U2OS cells when evaluated using FABCON^60^, a contact site reporter (Fig. S5C). These findings indicate that OPA1 does not function as a tether between LDs and mitochondria; instead, the data suggest a dynamic targeting mechanism allowing OPA1 to alternatively distribute between LDs and mitochondria.

### The naturally occurring OPA1 isoform 2, lacking exon 4, localizes exclusively to mitochondria

Splicing events at OPA1’s exon 4 have been documented in NCBI and UniProt databases. OPA1 isoform 2 (OPA1^iso2^) is a naturally occurring variant in which exon 4 is spliced out from the transcript (Fig. 4A). By analyzing the inclusion and exclusion of OPA1 exon 4 from the Genotype-Tissue Expression (GTEx) portal as an indicative expression readout of isoforms 1 and 2, respectively, we found that OPA1^iso2^ mRNA was widely expressed in most tissues, albeit at lower levels than OPA1^iso1^ mRNA (Fig. S6A). OPA1^iso2^ transcripts were also expressed in U2OS and HeLa cells (Fig. S6B). Given the importance of exon 4 in OPA1’s LD targeting, we decided to examine the subcellular distribution of OPA1^iso2^. We found YFP-tagged OPA1^iso2^ was only localized in mitochondria in OA-treated WT U2OS cells (Fig. 5A), sharply contrasting OPA1 ^iso1^’s distribution to LDs (Fig. 2A,B). Even with wMTS, which compromised mitochondrial import and enhanced OPA1 enrichment on LDs (Fig. 3E), the exon 4–omitting OPA1^iso2-wMTS^ remained majoritively localized to mitochondria in both U2OS and HeLa cells (Figs. 5A,B, and S6C,D).

**Figure 5.**
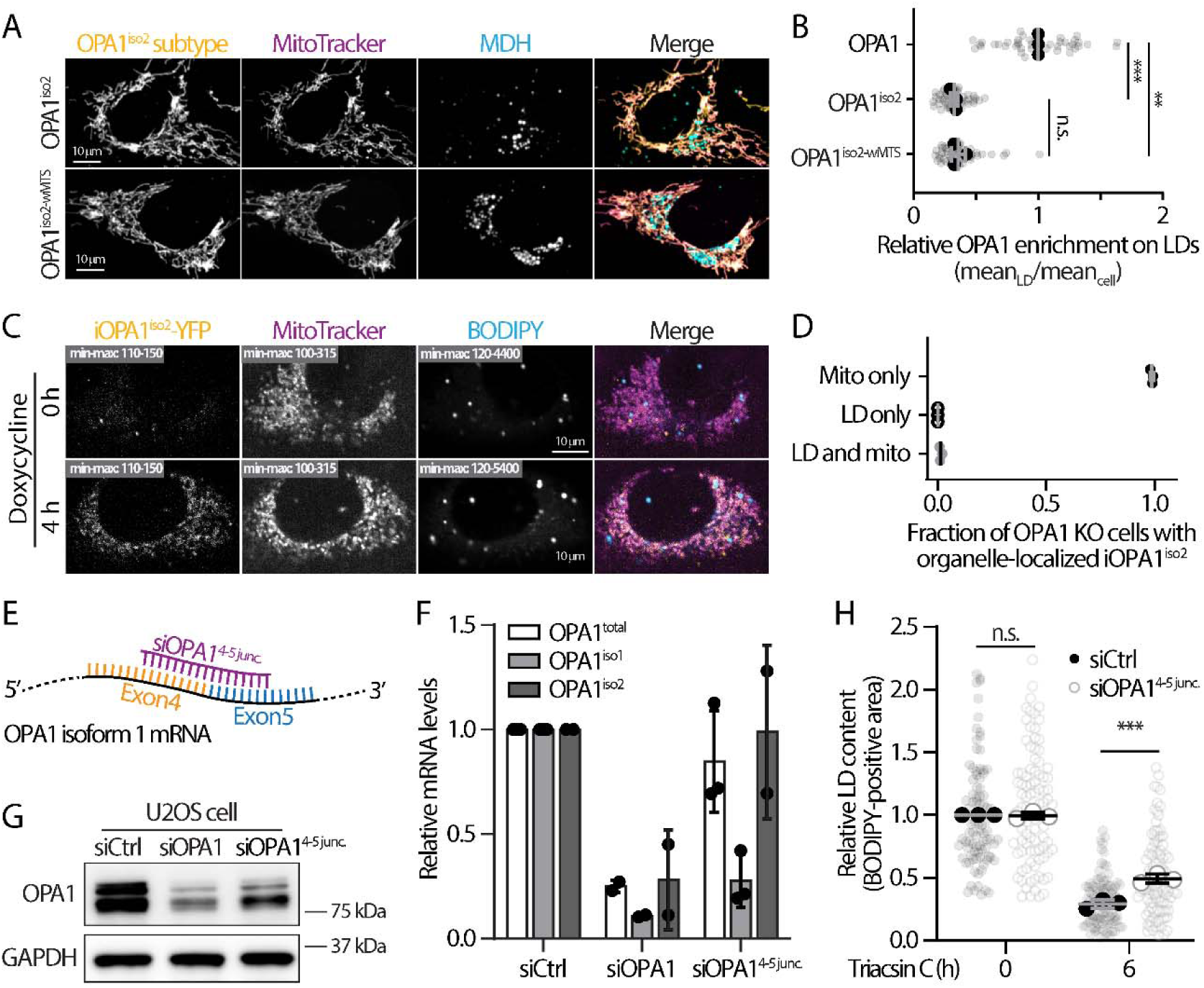
OPA1 isoform 2 lacks exon 4 and shows minimal localization to lipid droplets. **(A)** Localization of OPA1^iso2^-YFP and OPA1^iso2-wMTS^-YFP, mitochondria (MitoTracker Deep Red), lipid droplets (LDs; MDH) in U2OS cells treated with 100 µM oleic acid (OA) overnight detected by confocal microscopy. Maximal intensity projected confocal images from four axial slices (∼1 µm total thickness) are shown. **(B)** Relative enrichment of OPA1^iso2^ and OPA1^iso2-wMTS^ as described in (A) and OPA1 on LDs in U2OS cells treated with 100 µM OA overnight. Means ± standard deviation from three independent experiments are shown (total of 48–52 cells). n.s., no significance, ***P ≤ 0.001, **P ≤ 0.01, as assessed by one-way ANOVA. **(C)** Subcellular distribution of inducible OPA1^iso2^ (iOPA1 ^iso2^)-YFP on LDs (BODIPY 665/676) and mitochondria (MitoTracker Red) in *OPA1* knockout (KO) U2OS cells following 4 µg/mL doxycycline treatment for 4 h. Confocal images from a single axial slice are shown. **(D)** Fraction of cells with iOPA1^iso2^ localized to indicated organelle as described in (C). Means ± standard deviation from three independent experiments are shown. **(E)** Design of an siRNA targeting the OPA1 exon 4–exon 5 junction for selective depletion of isoform 1. **(F and G)** Levels of (F) OPA1 isoform-specific mRNA or (G) protein in U2OS cells transfected with the indicated siRNAs. Means ± standard deviation from two or three independent experiments are shown in (F). **(H)** Relative LD content in siCtrl- and siOPA1^4-5^ ^junc^-transfected U2OS cells treated with 100 µM OA overnight followed by 10 µM triacsin C incubation for 6 h. Means ± standard deviation from three independent experiments are shown (total of 90–101 cells). n.s., no significance, ***P ≤ 0.001, as assessed by one-way ANOVA.

Coincidently, when expressed in *OPA1* KO U2OS cells, inducible OPA1^iso2^ (iOPA1^iso2^-YFP) exclusively entered mitochondria with virtually no cells displaying any iOPA1^iso2^-YFP distributed to LDs (Fig. 5C,D). These observations further support our hypothesis that exon 4 is required for OPA1’s LD targeting.

We next examined whether OPA1^iso1^ alone is required for FA release from LDs using a siRNA targeting the exon 4–exon 5 junction (siOPA1^4-5^ ^junc^) (Fig. 5E). U2OS cells transfected with siOPA1^4-5^ ^junc^ showed a specific, substantial reduction of OPA1^iso1^ mRNA accompanied with a prominent depletion of L-OPA1, while a broader depletion of OPA1 mRNA and protein was observed in siOPA1-transfected cells (Fig. 5F,G). Intriguingly, siOPA1^4-5^ ^junc^-treated U2OS cells displayed significantly more accumulated LDs (49% remaining) than siControl (siCtrl)-transfected cells (29% remaining) after triacsin C treatment (Fig. 5H), which phenocopied the FA release defect observed in *OPA1* KO U2OS cells (Fig. 1C). A similar difference in LD content between siCtrl (34% remaining), siOPA1 (57% remaining), and siOPA1^4-5^ ^junc^ (53% remaining) following triacsin C treatment was also observed in HeLa cells (Fig. S6E). These results suggest that OPA1^iso1^ is required for FA release from LDs, likely through its ability to target LDs.

### LD-localized OPA1 facilitates recruitment of lipases

To gain further insights into how OPA1 mediates FA release from LDs, we interrogated whether OPA1 could facilitate lipases’ access to LDs, which is essential to lipase functions^3, 4^. Subcellular fractionation revealed that endogenous ATGL and HSL were significantly enriched in the LD fraction from WT U2OS cells compared to the fraction from *OPA1* KO U2OS cells (Fig. 6A,B).

**Figure. 6.**
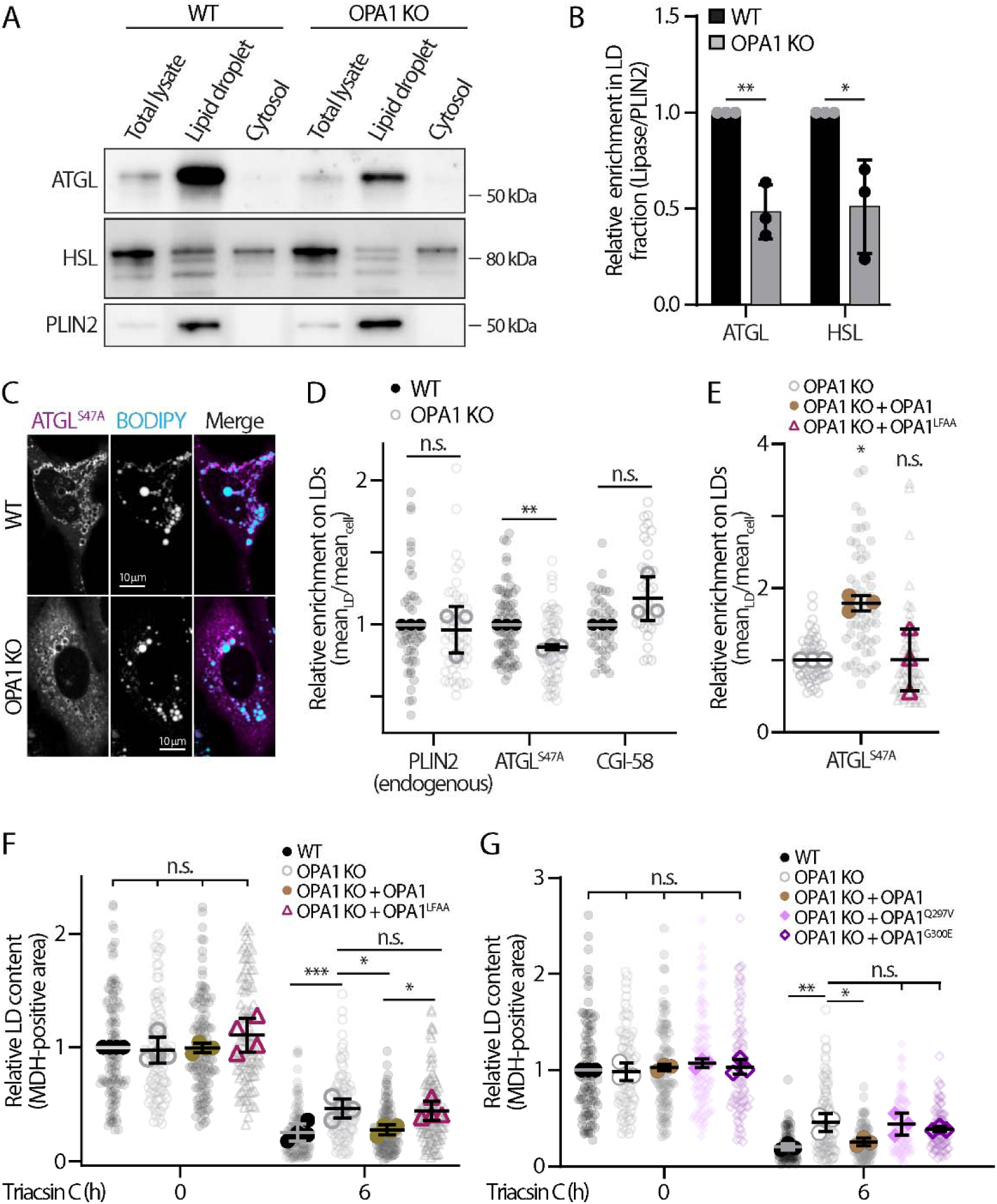
Lipid droplet–associated OPA1 facilitates lipase recruitment. **(A)** Western blot analysis of endogenous ATGL, HSL, and PLIN2 in sucrose-gradient cellular fractionations from wildtype (WT) and *OPA1* knockout (KO) U2OS cells treated with 500 μM oleic acid (OA). **(B)** ATGL and HSL abundance in lipid droplet (LD) fractions normalized to PLIN2 as described in (A). Means ± standard deviation from three independent experiments are shown. **P ≤ 0.01, *P ≤ 0.05, as assessed by unpaired *t*-test. **(C)** Localization of ATGL S47A-Halo and LDs (BODIPY-493/503) in WT and *OPA1* KO U2OS cells treated with 100 µM OA overnight. Maximal intensity projected confocal images from four axial slices (∼1 µm total thickness) are shown. **(D)** Relative enrichment of endogenous PLIN2 (detected by immunostaining), ATGL^S47A^-Halo, and mApple-CGI-58 on LDs from Figs. S5A, 5C, and S5B, respectively. Means ± standard deviation from three independent experiments are shown (n = 72–74 cells/experiment for ATGL^S47A^-Halo; n = 50–52 cells/experiment for PLIN2, and n = 34–46 cells/experiment for mApple-CGI-58). n.s., no significance, **P ≤ 0.01, as assessed by unpaired *t*-test. **(E)** Relative enrichment of ATGL^S47A^-Halo on LDs in *OPA1* KO U2OS cells and *OPA1* KO U2OS cells reconstituted with lenti-OPA1-YFP or lenti-OPA1^LFAA^-YFP. Means ± standard deviation from three independent experiments are shown (total of 56–60 cells). For statistics in panels E and F, n.s., no significance, **P ≤ 0.01, *P ≤ 0.05, as assessed by one-way ANOVA. **(F)** MDH-positive LD content in WT and *OPA1* KO U2OS cells, as well as *OPA1* KO U2OS cells reconstituted with lenti-OPA1-YFP or lenti-OPA1^LFAA^-YFP. Cells were treated with 100 µM OA overnight followed by 6-h incubation with 10 µM triacsin C. Meane ± standard deviation from four independent experiments are shown (total of 108–131 cells). (**G**) MDH-positive LD content in wildtype (WT) and *OPA1* KO U2OS cells, as well as *OPA1* KO cells reconstituted with lenti-OPA1-YFP, lenti-OPA1^Q297V^-YFP, or lenti-OPA1^G300E^-YFP. Cells were treated with 100 µM OA overnight followed by 6-h incubation with 10 µM triacsin C. Means ± standard deviation from three independent experiments are shown (total of 83–109 cells).

Consistent with endogenous ATGL, the enrichment of exogenous ATGL^S47A^-Halo, a catalytically dead ATGL used to assess localization without consuming LDs, was also significantly lower in *OPA1* KO cells than WT cells (Fig. 6C,D). Endogenous PLIN2, examined by fractionation and immunostaining, and overexpressed mApple-CGI-58, visualized by confocal microscopy, had comparable LD distribution across WT and *OPA1* KO U2OS cells (Figs. 6A,D and S7A,B), eliminating the possibility of a global reduction of the LD proteome in *OPA1* KO cells.

Reconstituting OPA1-YFP, but not OPA1^LFAA^-YFP with reduced LD enrichment, reversed the defects in ATGL^S47A^-Halo enrichment on LDs in *OPA1* KO U2OS cells (Fig. 6E). Similarly, OPA1-YFP restored defective FA release following triacsin C treatment in *OPA1* KO U2OS cells whereas OPA1^LFAA^-YFP had minimal effects (Fig. 6F). These findings indicate that OPA1 abundance on LDs is crucial for enriching ATGL and HSL before FAs are released from LDs.

OPA1’s GTPase activity is essential for its role in mediating mitochondria inner membrane fusion^61^, and we found this activity was also required for OPA1’s facilitation of FA release from LDs, as the two GTPase mutants, G300E and Q297V, which are defective in rescue mitochondrial fusion deficiency^62–65^, failed to rescue the FA release defect observed in *OPA1* KO cells (Fig. 6G). Given that the cellular expression and LD enrichment of CGI-58 was not changed due to OPA1 depletion (Figs. 6D, S1E, and S7B), we propose that OPA1-mediated lipase recruitment to LDs is independent of canonical lipolytic signaling.

### OPA1’s subcellular distribution differentially regulates mitochondrial characteristics

In addition to facilitating FA release, OPA1’s LD distribution competes with its mitochondrial targeting, likely reducing OPA1’s ability to enter mitochondria. When measuring OPA1’s relative mitochondrial enrichment, we observed a significant reduction of OPA1 present in mitochondria in OA-treated (LD-laden) U2OS cells compared to untreated cells (Fig. 7A); OPA1’s mitochondrial enrichment was negatively correlated with its enrichment on LDs (Fig. 2B).

**Figure 7.**
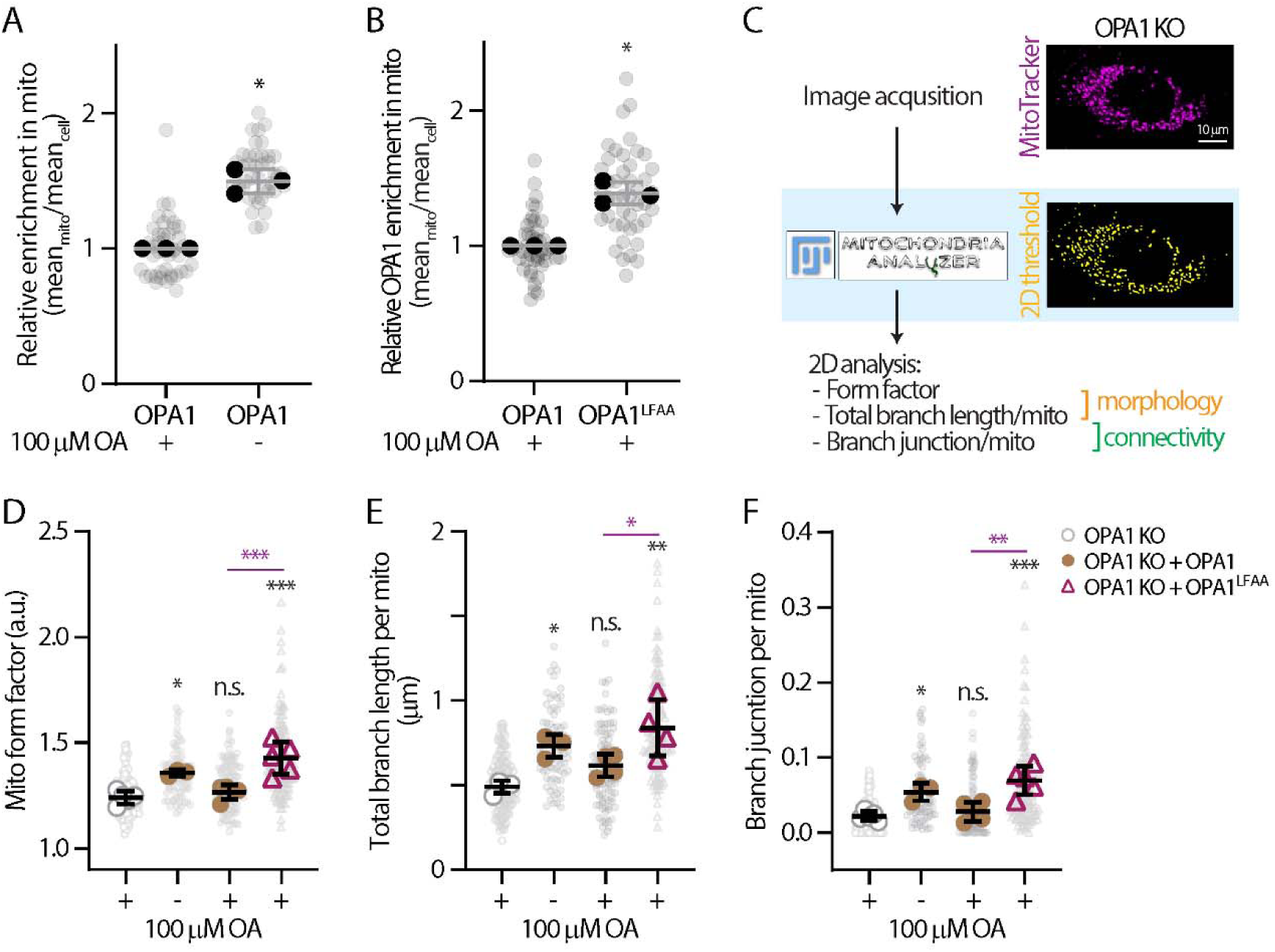
Differential regulation of mitochondrial characteristics depending on OPA1’s subcellular distribution. (A) Relative enrichment of OPA1-YFP in mitochondria in U2OS cells ± overnight 100 µM oleic acid (OA) treatment. Means ± standard deviation from three independent experiments are shown (total of 40–43 cells). *P ≤ 0.05, as assessed by paired *t*-test. **(B)** Relative enrichment of OPA1-YFP or OPA1^LFAA^-YFP in mitochondria in U2OS cells treated with overnight 100 µM OA. Means ± standard deviation from three independent experiments are shown (n = 45 cells/condition). **(C)** A schematic illustrating 2D analysis of mitochondrial morphology and connectivity using the Fiji plugin, ‘Mitochondria-Analyzer’. Representative MitoTracker DeepRed images (maximal intensity projected confocal images from three axial slices; 0.6 µm in thickness) from *OPA1* knockout (KO) U2OS cells and corresponding analyses are shown. **(D–F)** 2D analysis of mitochondrial morphology and connectivity measuring (D) form factor, (E) total branch length, and (F) branch junction in MitoTracker DeepRed-stained *OPA1* KO U2OS cells and *OPA1* KO U2OS cells reconstituted with lenti-OPA1-YFP or lenti-OPA1^LFAA^-YFP. Means ± standard deviation from three or four independent experiments are shown (total of 80–133 cells). n.s., no significance, ***P ≤ 0.001, **P ≤ 0.01, *P ≤ 0.05, as assessed by one-way ANOVA. Black asterisks represent comparisons to *OPA1* KO, and purple asterisks represent the indicated comparison.

Similarly, OPA1^LFAA^, which exhibited reduced association to LDs (Fig. 4J), had higher enrichment in mitochondria than WT OPA1 (Fig. 7B), further supporting the idea of competitive, alternative targeting between LDs and mitochondria. Given OPA1’s critical role in regulating mitochondrial inner membrane fusion, we reasoned that the reduced presence of OPA1 in mitochondria due to its partitioning to LDs might influence mitochondrial morphology. To test this idea, we used Mitochondria-Analyzer^66^, a Fiji plugin (Fig. 7C), to establish baseline 2D measurements of mitochondrial form factor, total branch length, and branch junctions as characteristics of morphology and connectivity in WT and *OPA1* KO U2OS cells. Consistent with prior studies^23, 67, 68^, *OPA1* KO cells had significantly lower mitochondrial form factor, shorter total branch length, and fewer branch junctions than the WT cells (Fig. S8A–D). While expressing exogenous OPA1-YFP in *OPA1* KO cells partially rescued mitochondrial morphology in untreated cells (Fig. 7D–F), OPA1-YFP failed to rescue mitochondrial characteristics in OA-treated *OPA1* KO cells, a condition under which effective mitochondrial entry of OPA1-YFP is reduced due to its LD distribution. Intriguingly, OPA1^LFAA^-YFP, with a reduced affinity for LDs, restored mitochondrial morphology and connectivity in OA-treated *OPA1* KO cells to levels similar to those observed in untreated OPA1-YFP–expressing *OPA1* KO cells (Fig. 7D–F). Together these results suggest the OPA1–LD interaction can dynamically regulate mitochondrial characteristics by modulating effective OPA1 entry into mitochondria.

### Effects of OPA1 S158N polymorphism on FA release, mitochondrial characteristics, and human metabolic traits

The most common polymorphism in the OPA1 coding sequence, a substitution of serine to asparagine at position 158 (p.Ser158Asn; c.473G>A, rs7624750), is located within exon 4 and has an average allele frequency of 0.4710 (gnomAD, Table S1). This substitution is accompanied by a relaxation of the N-terminal of the amphipathic helix as predicted by AlphaFold 3 (Fig. 8A), which further separates leucine^1^^56^ from the rest of the non-polar residues in exon 4, possibly leading to weakened LD binding. We observed a significant reduction in the relative enrichment of OPA1^S158N^-YFP on LDs and an increased distribution in mitochondria (Fig. 8B,C), mimicking the defective LD interactions of OPA1^LFAA^ with synthetic mutations. This led us to speculate if OPA1^S158N^ was also functionally distinct from OPA1 in FA release and mitochondrial characteristics. When repeating the OA withdrawal experiment, OPA1^S158N^ failed to rescue defects observed in FA release from LDs following triacsin C treatment in *OPA1* KO U2OS cells (Fig. 8D). However, OPA1^S158N^ increased mitochondrial form factor, total branch length, and branch junction regardless of the presence of LDs, contrasting the minimal rescue OPA1 provided in OA-treated *OPA1* KO cells (Fig. 8E–G). As OPA1^S158N^ is a natural variant and OPA1^LFAA^ is a synthetic mutant, these observations further strengthened the idea that exon 4-mediated dynamic distribution of OPA1 on LDs plays a crucial role in governing the release of FAs from LDs and regulating mitochondrial morphology and connectivity. Overall, these findings suggest OPA1 dynamically integrates the functions of both LDs and mitochondria via alternative organelle targeting.

**Figure 8.**
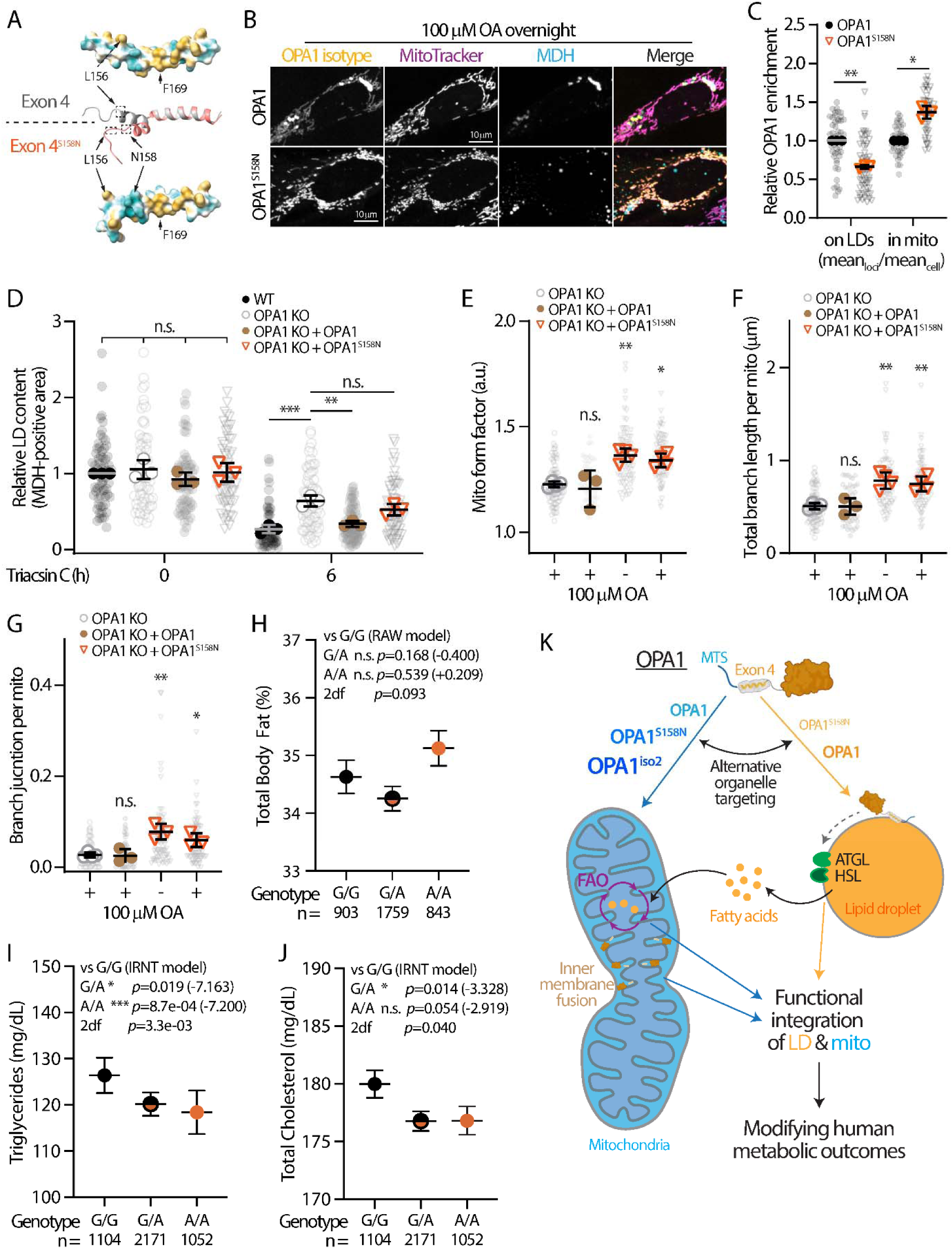
The S158N polymorphism within exon 4 impairs OPA1 targeting to lipid droplets and is associated with clinical manifestations in patients. (A) Amphipathic helix structures and hydrophobicity distributions of exon 4 and exon 4 S158N, predicted by AlphaFold3 and illustrated in ChimeraX. Cyan and yellow indicate polar and non-polar amino acids, respectively, and arrows denote locations of indicated residues. (B) Localization of YFP-tagged OPA1 and OPA1^S158N^ (lentiviral), mitochondria (MitoTracker Deep Red), and lipid droplets (LDs; MDH) in U2OS cells treated with 100 µM oleic acid (OA) overnight. Maximal intensity projected confocal images from four axial slices (∼1 µm total thickness) are shown. (C) Relative LD and mitochondria (mito) enrichment of OPA1 and OPA1^S158N^ in U2OS cells as described in (B). Means ± standard deviation from three independent experiments are shown (n = 45 cells/condition). **P ≤ 0.01, *P ≤ 0.05, as assessed by paired *t*-test. **(D)** MDH-positive LD content in wildtype (WT) and *OPA1* knockout (KO) U2OS cells, and in *OPA1* KO cells reconstituted with lenti-OPA1-YFP or lenti-OPA1^S158N^-YFP. Cells were treated with 100 µM OA overnight followed by 6-h incubation with 10 µM triacsin C. Means ± standard deviation from three independent experiments are shown (total of 63–83 cells). **(E–G)** Two-dimensional analysis of mitochondrial morphology and connectivity via measuring (E) form factor, (F) total branch length, and (G) branch junction in MitoTracker DeepRed-stained *OPA1* KO U2OS cells and *OPA1* KO cells reconstituted with lenti-OPA1-YFP or lenti-OPA1^S158N^-YFP. Means ± standard deviation from three or four independent experiments are shown (total of 55–83 cells). For panels D–G, n.s., no significance, ***P ≤ 0.001, **P ≤ 0.01, *P ≤ 0.05, as assessed by one-way ANOVA. **(H–J)** Clinical data was mined from the St. Jude LIFE database to assess (H) body fat percentage, (I) blood triacylglycerol levels, and (J) blood cholesterol levels in patients with different S158N genotypes. P-values and differences (within parentheses) for comparison of G/A or A/A vs G/G are indicated, along with p-values from the two-degrees-of-freedom model analysis. Body fat was analyzed using the raw value (RAW model), while triacylglycerol and cholesterol were analyzed using inverse rank-normal transformed values (IRNT model). **(K)** Proposed model of OPA1’s functions and impacts on LDs and mitochondria depending on its alternative targeting and its implication in human metabolic outcomes. OPA1-mediated lipase recruitment to LDs is indicated with a dashed arrow. Abbreviations: FAO, fatty acid oxidation; MTS, mitochondria targeting sequence.

It has already been established that functional coupling between LDs and mitochondria within cells can influence organismal metabolism^69, 70^. As serine^158^ and asparagine^158^ differentially regulate LD and mitochondrial functions, we speculated this polymorphism may also affect human metabolism. Using the St. Jude Lifetime Cohort Study^71^ of individuals with a history of childhood cancer diagnosed and treated at St. Jude Children’s Research Hospital between 1962 and 2012 who survived ≥5 years from diagnosis, we identified lipid-related metabolic traits strongly correlated with this polymorphism. While body fat content trended towards higher values in A/A individuals, this trend was not statistically significant (G/A vs G/G, P = 0.168; A/A vs G/G, P = 0.539; 2df, P = 0.093) (Fig. 8H and Table S2); therefore, we treated this as a suggestive, secondary observation rather than evidence of altered fat accumulation. Intriguingly, the strongest and most consistent signal was seen for blood TAGs, for which the A allele was associated with lower TAG levels (Fig. 8I and Table S2). In the codominant model, both G/A and A/A had lower TAG levels relative to G/G, with the A/A contrast the most significant (P = 8.7 × 10⁻ ; 2df, P = 3.3 × 10⁻^3^). Similarly, the A allele was associated with lower total cholesterol (Fig. 8J): G/A carriers had significantly lower total cholesterol than G/G carriers. A/A carriers had a trend similar to that observed with G/A carries, but it was not significantly different from G/G carriers. The association of the A allele with high-density lipoprotein (HDL) cholesterol was positive; A/A carriers had the highest level of HDL, while G/G carriers had the lowest, which is consistent with a potentially more favorable lipid profile overall (Table S2). The association of A allele with lower TAG and cholesterol was unexpected, as these metrics are all often linked to body fat content^72–75^. This dissociation between body fat and blood TAG and cholesterol suggests that OPA1 may exert pleiotropic effects through context-dependent differences in lipid droplet–mitochondrial integration across tissues and cell types. Although these asparagine^158^-associated metabolic traits were not clinically significant compared to the WT (consistent with the gnomAD database’s benign classification of this polymorphism), these findings support the theory that these polymorphisms have modest but detectable effects, with both exhibiting potentially beneficial and unfavorable associations depending on the phenotype. We emphasize that these human associations are modest and, for body fat, non-significant; they are consistent with, but not on their own sufficient to establish, an effect of S158N on adiposity. The most robust human evidence pertains to circulating lipids rather than fat accumulation. Altogether, these findings suggest that the differential coordination of FA release from LDs and mitochondrial dynamics by asparagine^158^ polymorphism is clinically relevant as a metabolic modifier in pediatric cancer survivors.

## Discussion

FAs are essential biomolecules playing critical roles in energy production, lipid homeostasis, and signal transduction. Nearly all cell types can store FAs and utilize these stockpiles when needed; our findings reveal novel mechanisms governing the utilization of these FA stockpiles (Fig. 8K). We first demonstrated that a subpopulation of OPA1 is localized to LDs through the exon 4 mediated targeting. When localized to LDs, OPA1 promotes the recruitment of lipases, facilitating the consequent FA release. This interaction dynamically controls OPA1’s entry into mitochondria, where OPA1 influences mitochondrial fusion and connectivity. We have established a mechanistic link between mitochondrial morphology and LD homeostasis. Finally, we revealed that OPA1’s alternative organelle distribution can be modulated by a common polymorphism in exon 4; this may be relevant to human metabolism, as circulating lipid traits were associated with this polymorphism in pediatric cancer survivors, whereas the trend toward higher body fat was not significant. Overall, these findings broaden our collective understanding of OPA1’s role in lipid homeostasis and organelle crosstalk, providing new insights on the dynamic interplay between FA storage and mitochondrial morphology. Recent discoveries revealing non-canonical roles for OPA1 in stress responses, ferroptosis, and transcriptional activation within antitumor immunity^76, 77^ emphasize the growing need for a deeper mechanistic understanding of this protein. This timely study provides important insights that help explain these emerging findings.

Lipolysis has been extensively studied in adipose tissues, the primary body fat storage depot in animals, and PKA plays an essential role in this process via the inducible recruitment of ATGL and HSL to LDs through a series of phosphorylation events. While emerging evidence indicates the existence of lipolysis in a variety of non-adipose cell types^4, 15^, this canonical view of lipolysis becomes insufficient because PLIN1, one of the key effectors downstream of PKA, is only expressed in selective tissues while lipases are widely expressed. As OPA1 is also broadly expressed in many human tissues and OPA1-mediated FA release occurs independently of canonical lipolytic signaling, we propose a new explanation for how non-adipose cell types dynamically regulate and utilize FAs within LDs.

The mechanism behind OPA1’s ability to facilitate lipase recruitment is a question that warrants further investigation, as ATGL and HSL’s association with LDs is subject to complex regulations. In addition to lipolytic stimulation that dynamically recruits and activates ATGL on LDs via interactions with CGI-58 in adipocytes^46–48^, ATGL also binds LDs directly via a C-terminal hydrophobic domain^46, 78, 79^ and can be delivered to LDs via Arf1/COPI machinery^80–82^. Similarly, HSL recruitment to LDs is regulated by both PKA-dependent phosphorylation and PKA-independent LD-binding motifs^83, 84^. Beyond these mechanisms, LD’s physical properties, such as their size and levels of lipid packing defects, can broadly influence protein targeting^85, 86^. OPA1 may regulate ATGL and HSL recruitment through any one of these mechanisms. Notably, we observed that both endogenous and exogenous OPA1 frequently localize on clusters of small LDs, which have been characterized by a higher surface-to-volume ratio than large LDs and are associated with enhanced lipolytic activity^87^. Given OPA1’s established role in membrane remodeling via its GTPase activity ^88, 89^, our finding of the requirement of OPA1’s GTPase activity in supporting FA release from LDs suggests that OPA1 enhances lipase recruitment or activation by altering the physical nature of LDs. Membrane-morphing endosomal sorting complexes required for transport (ESCRT) components have been implied to play a role in facilitating inter-organelle FA trafficking^90, 91^, lending further support to this hypothesis. Therefore, it will be important to determine how LD-associated OPA1 that possesses membrane-remodeling capabilities contributes to lipase recruitment and FA release.

The data presented here may begin to resolve previous conflicting observations^16–22^ regarding mitochondrial morphology and LD abundance under distinct metabolic conditions and perturbations from a technical point of view. OPA1’s role in mediating FA release may have been previously unrecognized due to the technical challenges related to the study of the tightly coupled functional interactions between LDs and mitochondria. By developing new FA flux assays with acute pharmacological perturbations designed to specifically assess trafficking routes across LDs and mitochondria, we found that OPA1 actively regulates FA release from LDs and mitochondrial FAO while minimally influencing FA import into mitochondria. This indicates FA trafficking routes between organelles are not directly coupled and emphasizes the importance of rigorously dissecting individual pathways to untangle the specific mechanisms underlying mitochondrial morphology and LD metabolism. Additional factors, such as heterogeneity of metabolically distinct mitochondria^92^ and mitochondria-mediated lipogenesis^93^, should be considered and examined thoroughly to achieve a comprehensive understanding of these processes.

To gain further insights into the regulation of OPA1 alternative organelle targeting, it is important to interrogate the mechanisms underlying the exon 4 splicing event. Our current analysis from the GTEx database revealed that heart and skeletal muscle have a higher inclusion of exon 4, whereas adipose tissue has a lower inclusion. Because heart and skeletal muscle primarily rely on FAO for ATP production^94, 95^, these data suggest a potential correlation between exon 4 inclusion and FA utilization. It is still unknown how OPA1’s exon 4 (or OPA1 itself) senses metabolic cues within the cytoplasm to determine its organelle distribution. Both pyruvate and FAs drive the TCA cycle via the acetyl-CoA intermediate, raising the possibility that OPA1 may respond to these metabolites and dynamically adjust its organelle distribution based on the sensed fuel sources. Studying polymorphisms within exon 4 may provide additional clues regarding the metabolic regulation in OPA1’s subcellular localization. According to the gnomAD database, several polymorphisms with low allele frequencies, such as p.Ser177Ile and p.Ile152Asn (Table S1), may also alter the non-polar interface of exon 4’s amphipathic helix and potentially affect OPA1’s interactions with LDs.

In addition, given that OPA1’s localization to LDs is determined by the competition between exon 4 and OPA1’s MTS, it is important to understand whether OPA1’s mitochondrial translocation itself can be regulated. While our computational prediction suggested that OPA1 has a strong MTS, a previous study identified phosphorylation events at serine^21^ and ^22^ in OPA1’s N-terminal region^96^, which could modulate MTS strength. Furthermore, the MitoFates prediction algorithm^58^ suggests OPA1’s first 39 amino acids could function as a weak MTS, with a probability score of 0.509 (where 1 represents maximum confidence). Interestingly, similarly low MTS probabilities have been associated with the inefficient mitochondrial import of ATFS-1 in *Caenorhabditis elegans*^97^. Given the functional importance of OPA1 at both LDs and mitochondria, further studies illuminating the mechanisms by which OPA1 differentially targets to organelles should be prioritized. Our current data on the asparagine^158^ polymorphism (A allele) implicates the importance of OPA1’s alternative organelle targeting via a functional trade-off and the integration between LDs and mitochondria. This trade-off likely underlies the differentia metabolic traits associated with serine^158^ and asparagine^158^, as each provides specific metabolic benefits. Integrating experimental interrogation of OPA1’s natural polymorphisms, along with continued MTS interrogation and clinical data mining, will likely uncover additional roles for OPA1 in human health and physiology through its complex moderation of LDs, mitochondria, and their interactions under various cell states.

Together, these findings highlight OPA1 as a previously unrecognized coordinator for LD–mitochondria communication via alternative organelle targeting and provide an orthogonal view of the functional coupling between LD and mitochondria beyond physical contact sites. We propose that OPA1 acts as a molecular integrator aligning mitochondrial dynamics with lipid storage and utilization, enabling energy adaptation in response to cellular physiological demands.

## Materials and Methods

### Cell lines, transfection, and transduction

U2OS cells (HTB-96), HeLa (CCL-2) cells, and *OPA1* KO MEFs (CRL-2995) were purchased from American Type Culture Collection (ATCC); Huh7 (JCRB0403) cells were purchased from the Japanese Cancer Research Bank. The WT MEFs were generated previously^98, 99^. All cells were maintained in their respective base media supplemented with 10% fetal bovine serum (FBS) and 1X penicillin/streptomycin solution per vendors’ recommendation. The base media used were McCoy’s 5A (Cat# 16600082, Thermo Fisher Scientific) for U2OS cells, Eagle’s Minimal Essential Medium (EMEM) (Cat# 30-2003, ATCC) for HeLa cells, and high-glucose Dulbecco’s Modified Eagle Medium (DMEM) (Cat# 11995-065, Thermo Fisher Scientific) for Huh7 cells and MEFs. Cells were routinely tested via the MycoAlert® Plus Mycoplasma Detection kit and found free of mycoplasma contamination (Cat# LT07-703, Lonza).

Transfection and transduction were performed 16–24 h after cell seeding. DNA plasmid transfection was performed using TransIT-LT1 (Muris Bio LLC) for HeLa cells and Lipofectamine 3000 (Thermo Fisher Scientific) for U2OS and Huh7 cells with 50–500 ng DNA for 16–20 h according to manufacturer’s instructions. To knock down OPA1, HeLa or U2OS cells were transfected with 25 nM scramble (Cat# D-001810-10), OPA1 siRNA (Cat# L-005273-00-0020), or custom designed siRNA targeting OPA1’s exon 4–5 junction (sense: 5’ CCUCAGGUUCUCCGGAAGAUU 3’, antisense: 5’ UCUUCCGGAGAACCU GAGGUU 3’ (Dharmacon) using TransIT-TKO (Mirus Bio LLC) according to the manufacturer’s instructions for 48 h. Lentiviral particles were transduced into cells in the presence of 10 μg/mL protamine sulfate 40–48 h prior to the experiment. Multiplicity of infection (MOI) used were: 5 for TOM20-Halo, 20–100 for OPA1-YFP isoforms and mutants, 50 for OPA1-mhYFP, and 100 for iOPA1-YFP isoforms.

### Generation of OPA1 KO U2OS cells

OPA1-/-U2OS cells were generated using CRISPR technology in the Center for Advanced Genome Engineering at St. Jude Children’s Research Hospital [St. Jude]. First, 500,000 U2OS cells were transiently transfected with precomplexed ribonuclear proteins (RNPs) consisting of 150 pmol of chemically modified sgRNA (CAGE2484.OPA1.g3 spacer – 5’-CCUAUUUAAAG AUAGUUCUC -3’, IDT) and 50 pmol of 3X NLS SpCas9 protein (St. Jude Protein Production Core). Then, 300 ng of pMaxGFP (Lonza) plasmid was added to the transfection mix and transfected via nucleofection (Lonza, 4D-Nucleofector™ X-unit) according to the manufacturer’s protocol using solution P3 and program CM-104 in a 20-μL cuvette. Five days post nucleofection, cells were single-cell sorted by flow cytometry at the St. Jude Flow Cytometry and Cell Sorting Shared Resource Center to enrich for GFP+ (transfected) cells into 96-well tissue culture treated plates containing pre-warmed McCoy’s 5A media. Cells were clonally expanded and screened for the desired modification (out-of-frame indels) via targeted amplicon sequencing using gene specific primers with partial Illumina adapter overhangs (CAGE2484.OPA1.DS.F– 5’-CTACACGACGCTCTTCCGATCTtggatttgtgcaat gcagtagccct-3’ and CAGE2484.OPA1.DS.R-5’-CAGACGTGTGCTCTTCCGATCT-ccttacctcagggctaacggtacagc-3’, overhangs shown in uppercase) as previously described^100^. Next-generation sequencing analysis of clones was performed using CRIS.py^101^. Final clones were authenticated using the PowerPlex® Fusion System (Promega) performed by the Hartwell Center for Biotechnology at St. Jude. Final clones were tested using the MycoAlertTMPlus Mycoplasma Detection Kit (Lonza) and found to be negative for mycoplasma. U2OS cells were further validated by western blotting and immunostaining.

### Reagents

BODIPY 493/503 (Cat# D3922), BODIPY 665/676 (Cat# B3932), sodium pyruvate (Cat# 11360070), D-glucose (Cat# A2494001), trypsin-EDTA (Cat# 25200056), PBS (Cat# 10010023), FBS (Cat# A5669801), Hoechst (Cat# 62249), MitoTracker Red CMXRos (Cat# M7512), and MitoTracker Deep Red FM (Cat# M22426) were purchased from Thermo Fisher Scientific. AUTODOT^TM^ Visualization Dye, monodansylpentane (MDH; Cat# SM1000a) was purchased from Abcepta, oleic acid–BSA complex, referred to as oleic acid (OA), was purchased from Sigma-Aldrich (Cat# O3008) and Cayman Chemicals (Cat# 29557). TopFluor-OA (Cat# 810259) and NBD-arachidonic acid (Cat# 810106) were purchased from Avanti Research. BODIPY-linoleic acid (Cat# 29795), DGAT1 inhibitor (PF-04620110, Cat# 16425), DGAT2 inhibitor (PF-06424439, Cat# 17680), and triacsin C (Cat# 10007448) were purchased from Cayman Chemical. NG-497 (Cat# SML3712-5MG), Lalistat-2 (Cat# SML2053-5MG), and 2-deoxy-D-glucose (Cat# D8375-10MG) were purchased from Sigma-Aldrich. Doxycycline was purchased from Takara (Cat# 631311). HaloTag ligand JF646 (Cat# HT1060) was purchased from Promega.

### DNA plasmids

OPA1-YFP was generated by inserting the *OPA1* fragment amplified from U2OS cDNA into the pEYFP-N1 (Takara) plasmid between the *Sal*I and *Bam*HI sites. OPA1-YFP lentiviral plasmid (lenti-OPA1-YFP) was generated by inserting the OPA1-YFP fragment cut from the OPA1-YFP plasmid into the SJL12-CMV-GFP lentiviral backbone (provided by the St. Jude Vector Laboratory) between the *Xho*I and *Not*I restriction sites, followed by the removal of the Kozak sequence immediately 5’ of the YFP by site-directed mutagenesis. OPA1^wMTS^-YFP was generated by cloning a gBlock substituting all but two positively charged residues within OPA1’s MTS with alanine (as shown in Fig. 3C) into the SJL-CMV-OPA1-YFP between the *Xho*I and *Cla*I restriction sites, replacing the WT OPA1 MTS. OPA1^iso2^ and OPA1 mutants were generated by site-directed mutagenesis using lenti-OPA1-YFP as the template. To generate inducible OPA1-YFP lentiviral plasmids, a GFP-free TRE-mCh-E-GFP3G backbone (a Tet-on inducible plasmid provided by the St. Jude Vector Laboratory) was generated first by site-directed mutagenesis, followed by OPA1-YFP or OPA1^iso2^-YFP fragment insertion between the *Blp*I and *Bsi*WI sites using InFusion to create TRE-iOPA1-YFP-E3G and TRE-iOPA1^iso2^-YFP-E3G, respectively. YFP-tagged OPA1 exon4 (exon 4-YFP) was generated by inserting a gblock containing the exon 4 coding sequence (synthesized by IDT, Newark, NJ) into pEYFP-N1 between the *Sal*I and *Bam*HI restriction sites, followed by the removal of the Kozak sequence immediately 5’ of the YFP by site-directed mutagenesis. Exon 4-YFP mutants were generated by site-directed mutagenesis. Monomeric hyperfolder-YFP (mhYFP)-tagged OPA1 exon 4 was generated by replacing YFP in Exon 4-YFP with a mhYFP gBlock using *Bam*HI and *Bsr*GI restriction sites. OPA1-mhYFP was cloned by replacing YFP in the OPA1-YFP plasmid with a mhYFP gBlock using the *Age*I and *Bsr*GI restriction sites. Lenti-OPA1-mhYFP was subsequently generated by InFusion with an OPA1-mhYFP PCR fragment and a linearized SJL12-CMV-GFP backbone digested by *Xho*I and *Not*I restriction enzymes. Lenti-ATGL^S47A^-Halo was generated by inserting an ATGL PCR fragment amplified from U2OS cDNA and a Halo fragment into SJL12-CMV-GFP backbone using *Bsi*WI, *Age*I, and *Not*I restriction sites, followed by replacing serine^47^ with alanine by site-directed mutagenesis. To generate mApple-CGI-58, CGI-58 fragment was amplified from U2OS cDNA and inserted into the mApple-C1 backbone plasmid using the *Hind*III and *Bam*HI restriction sites. The pEGFP-C1-PLIN2 plasmid was purchased from Addgene (Plasmid #87161). The Lenti-TOM20-Halo plasmid was generated by Genscript (Piscataway, NJ, USA) using a pGenLenti backbone and a FLASH gene synthesis fragment containing a TOM20-Halo coding sequence from Halo-TOM20-N-10 (Addgene plasmid #123284). Production and titration of lentivirus were performed by St. Jude Vector Laboratory as described previously^60, 102^. Oligonucleotides used for cloning can be found in Table S2. All plasmids used in this work were confirmed by sequencing.

### Generation of cDNA, PCR, and real-time PCR

Total RNA from U2OS and HeLa cells was extracted using the PureLink^TM^ RNA Mini Kit (Thermo Fisher Scientific, Cat# 12183018A) according to the manufacturer’s instructions. Equal amounts of RNA (0.5–1 µg) were reverse-transcribed into cDNA using the SuperScript^TM^ IV First-Strand Synthesis System Kit (Thermo Fisher Scientific, Cat# 18091150) with oligo-dT primer.

Quantitative real-time PCR was performed on the QuantStudio 6 Pro Real-Time PCR System using the PowerUp™ SYBR™ Green Master Mix Kit (Thermo Fisher Scientific, Cat# A25918). Oligonucleotides used for real-time PCR are listed in Table S3.

### Metabolomics and lipidomic analysis

U2OS WT and *OPA1* KO cells were grown in a 15-cm dish to full confluency, scraped and harvested in cold PBS. Cell pellets were snap-frozen in liquid nitrogen and stored at -80°C before shipping to Metabolon (Morrisville, NC, USA) for metabolomic and lipidomic analysis. For metabolomics analysis, samples were prepared using the automated MicroLab STAR® system from Hamilton Company. Several recovery standards were added prior to the first step in the extraction process for quality control purposes. To remove protein, dissociate small molecules bound to protein or trapped in the precipitated protein matrix, and to recover chemically diverse metabolites, proteins were precipitated with methanol and vigorous shaking for 2 min (Glen Mills GenoGrinder 2000) followed by centrifugation at 500 g for 10 min at 12°C. The resulting extract was divided into equal fractions: two for analysis by two separate reverse phase (RP)/UPLC-MS/MS methods with positive ion mode electrospray ionization (ESI), one for analysis by RP/UPLC-MS/MS with negative ion mode ESI, one for analysis by HILIC/UPLC-MS/MS with negative ion mode ESI, while the remaining fractions were reserved for backup. Samples were placed briefly on a TurboVap® (Zymark) to remove the organic solvent. All methods utilized a Waters ACQUITY ultra-performance liquid chromatography (UPLC) and a Thermo Scientific Q-Exactive high resolution/accurate mass spectrometer interfaced with a heated electrospray ionization (HESI-II) source and Orbitrap mass analyzer operated at 35,000 mass resolution^103^. The dried sample extract was then reconstituted in solvents compatible to each of the following four methods. Each reconstitution solvent contained a series of standards at fixed concentrations to ensure injection and chromatographic consistency.

One (1) aliquot was analyzed using acidic positive ion conditions, chromatographically optimized for more hydrophilic compounds (PosEarly). In this method, the extract was gradient eluted from a C18 column (Waters UPLC BEH C18-2.1x100 mm, 1.7 µm) using water and methanol, containing 0.05% perfluoropentanoic acid (PFPA) and 0.1% formic acid. The second (2) aliquot was also analyzed using acidic positive ion conditions; however, it was chromatographically optimized for more hydrophobic compounds (PosLate). The extract was gradient eluted from the same aforementioned C18 column using methanol, acetonitrile, water, 0.05% PFPA, and 0.01% formic acid; the column was operated at an overall higher organic content. The third (3) aliquot was analyzed using basic negative ion optimized conditions using a separate dedicated C18 column (Neg). The basic extracts were gradient eluted from the column using methanol and water with 6.5 mM ammonium bicarbonate at pH 8. The fourth and final (4) aliquot was analyzed via negative ionization following elution from a hydrophilic interaction chromatography (HILIC) column (Waters UPLC BEH Amide 2.1x150 mm, 1.7 µm) using a gradient consisting of water and acetonitrile with 10 mM ammonium formate, pH 10.8. The MS analysis alternated between MS and data-dependent MSn scans using dynamic exclusion. The scan range varied slightly between methods but covered 70–1000 m/z. Raw data was extracted, peaks identified, and quality control was processed using a combination of Metabolon-developed software services (applications), and peaks were quantified using area-under-the-curve.

For lipidomics analysis, samples were homogenized in deionized water. About 11% of each homogenate was reserved for Bradford (protein) and/or DNA quantification for normalization purposes. The rest of each homogenate was subjected to a modified Bligh-Dyer extraction using methanol/water/dichloromethane in the presence of internal standards. The extracts were concentrated under nitrogen and reconstituted in 0.25 mL 10 mM ammonium acetate dichloromethane:methanol (50:50). The extracts were transferred to inserts and placed in vials for infusion-MS analysis, performed on a Shimazdu LC with nano PEEK tubing and the Sciex SelexIon-5500 QTRAP. The samples were analyzed via both positive and negative mode electrospray. The 5500 QTRAP scan was performed in multiple reaction monitoring (MRM) mode with the total of more than 1,100 MRMs. Individual lipid species were quantified by taking the peak area ratios of target compounds and their assigned internal standards, then multiplying by the concentration of internal standard added to the sample. Lipid species concentrations were background-subtracted using the concentrations detected in process blanks (water extracts) and run day normalized (when applicable). The resulting background-subtracted, run-day normalized lipid species concentrations were then used to calculate the lipid class and FA total concentrations, as well as the mol% composition values for lipid species, lipid classes, and FAs.

### FAO analysis

FAO in U2OS cells was measured as described previously^104^. WT (50,000 cells/well) or *OPA1* KO (75,000 cells/well) cells were cultured in a volume of 0.5 mL/well in a 24-well plate. Three days after seeding, cells were visually confirmed to have a confluent monolayer. To measure FAO, a prewarmed mixture of 1 mM L-carnitine, 0.3% BSA, 100 µM oleate, and 0.4 µCi ^14^C-oleate in serum-free McCoy’s 5A media with or without 5 µM Triacsin C was added to the cells. After a 3-h incubation at 37°C, 400 µL supernatant was added to 1.7-mL tubes for the collection of complete and incomplete FAO. Complete (^14^CO_2_ adsorbed on 1 M NaOH-soaked Whatman paper in cap) and incomplete FAO (soluble in 1 M perchloric acid) were measured with a Tri-Carb 2910TR liquid scintillation analyzer (Perkin-Elmer) for 3 min per vial in 3 mL Ultima Gold scintillation fluid (Sigma-Aldrich).

### Subcellular fractionation and western blotting

For subcellular fractionation, 6x 15-cm dish U2OS cells treated with 500 μM OA for 24 h at 100% confluency were harvested and lysed in ice-cold hypotonic lysis buffer (20 mM Tris-HC, pH 7.4, 1 mM EDTA, pH 8.0) supplemented with proteinase inhibitor cocktail via passing through 27G g needles 40 times. Mouse livers were lysed in the same lysis buffer using a Dounce homogenizer (tight pestle). Cellular fractions were isolated via ultracentrifugation at 50,000 x g for 1 h as previously described^60, 105^. Protein concentration of subcellular fractions and clear lysates were quantified using the Pierce BCA Protein Assay (Thermo Fisher Scientific, Cat# A65453). Equal amount of protein was denatured in 1x SDS-loading buffer at 98°C for 5– 10 min. This protocol provides purity and yield for LD but could rupture mitochondria in U2OS cells due to swelling in hypotonic buffer and/or high G-force spin.

To acquire intact mitochondria for proteinase K protection assay, U2OS cells were harvested via trypsinization and lysed in isotonic lysis buffer (20 mM Tris-HCl supplemented with 250 mM sucrose and 1 mM EDTA, pH 7.4) via a Dounce homogenizer (tight pestle) for 140–180 strokes until 70% of remaining cells were trypan-blue positive. Clear lysates were first collected by centrifugation of homogenates at 1,000 x g for 10 min at 4°C, followed by 20,000 x g centrifugation for 20 min at 4°C. Approximately 200 µL of the floating LD fraction was then carefully collected from the top of the cell lysate. LD fractions from mouse liver and U2OS cells (isotonic purification) were incubated with 10 µg/mL of proteinase K on ice for 2 min, followed by 2 mM phenylmethylsulfonyl fluoride deactivation for 3 min and denaturation in 1x SDS-loading buffer at 98°C for 10 min.

An equal amount of protein was loaded into an SDS-PAGE protein gel before being transferred to a nitrocellulose (Bio-Rad) or PVDF (Bio-Rad) membrane for western blotting. Membranes were blocked with EveryBlot blocking buffer (Bio-Rad) for 1 h before being incubated with a primary antibody at room temperature for 1 h or at 4°C for overnight. After incubation, the membranes were then washed with PBST for 10 min three times and incubated with a corresponding HRP conjugated secondary antibodies (1:3,000). The membranes were then washed again with PBST for 10 min three times before incubated with ECL substrates (Bio-Rad) and imaged with ChemiDoc (Bio-Rad) for chemiluminescence. Primary antibodies for western blotting were used at a 1:2000 dilution, which included anti-OPA1 antibody (Cat# 80471S, Cell Signaling Technology), anti-ADFP (PLIN2) antibody (Cat# AP5811c; Abcepta), anti-prohibitin 1(PHB1) antibody (Cat# PA5-19556, Invitrogen; Cat# 2426S, Cell Signaling Technology ), anti-ATGL antibody (Cat# 2439, Cell Signaling Technology), anti-HSL antibody (Cat# 4107T, Cell Signaling Technology), anti-CGI-58 antibody (Cat# 183739, Abcam), and anti-GAPDH antibody (Cat# 2118; Cell Signaling Technology). Secondary antibodies used were goat anti-rabbit IgG-HRP (Cat# 1706515, Bio-Rad) and goat anti-mouse IgG-HRP (Cat# 1706516, Bio-Rad).

### Immunostaining

Cells were fixed and immunostained as described previously^60^. All procedures were performed at room temperature while avoiding light, and all wash steps were using PBS unless otherwise indicated. First, cells were rinsed and fixed with 4% paraformaldehyde (PFA, Electron Microscopy Sciences) and 0.05% glutaraldehyde (GA, Electron Microscopy Sciences) in PBS for 15 min. Fixed cells were washed three times and permeabilized with 0.3% Triton X-100 in PBS for 20 min. Permeabilized cells were then blocked with 5% normal donkey or goat serum in PBS for 1 h followed by incubation with primary antibody (1:200 in dilution; anti-OPA1 antibody, Cat# 67589, Cell Signaling Technology; anti-PLIN2 (Cat# AP5118c, Abcepta); anti-TOM20 antibody (Cat# ab186735 or ab56783, Abcam) in PBS with 1% BSA (Sigma-Aldrich) at 4°C overnight. The next day, the samples were washed three times before being incubated with a secondary antibody, including Alexa Fluor Plus 488 conjugated donkey anti-rabbit IgG (Cat# A32790), Alexa Fluor Plus 555 conjugated goat anti-rabbit IgG (Cat# A32732), or Alexa Fluor Plus 647 conjugated goat anti-rabbit IgG (Cat# A32733, Thermo Fisher Scientific) for 1 h. The stained samples were washed three times and imaged using confocal microscopy.

### Fluorescence microscopy imaging, electron microscopy, and image analysis

All imaging experiments were performed on a custom-built Nikon microscope equipped with a Yokogawa CSU-W1 Spinning Disk unit, Z-axis piezo nanopositioners, and Tokai Hit STXG CO_2_ incubation system using a 60x (CFI APO TIRF, NA 1.49 oil) objective. Unless otherwise indicated, all cells were grown and transiently transfected with plasmids or transduced with lentiviral particles on #1.5 cover glasses imaging vessels, including Lab-Tek II chambered cover glasses (Cat# Z734853-96EA, Thermo Fisher Scientific) and MatTek dishes (Cat# P35G-1.5-10-C, MatTek). Prior to live-cell imaging, media were replaced with Fluoro-Brite DMEM (Thermo Fisher Scientific) supplemented with 5% FBS and penicillin/streptomycin and then imaged at 37°C supplemented with 7% CO_2_ in humidified air. Immunostaining samples were imaged in PBS at room temperature. All image analyses were performed with Fiji^106^ unless otherwise indicated. For intensity-based analysis, all intensity-based analyses were subjected to background (area without cells) subtraction.

#### OA withdrawa

To evaluate the FA release from LDs, cells seeded in 8-well chambered cover glass with transfection, transduction, and/or perturbation were treated with 100 µM OA overnight to induce LD formation. The next day, OA-containing media were replaced with fresh OA-free media with 10 μM triacsin C or without glucose to induce FA release from LDs. Cells were then stained with BODIPY-493/503 (500 ng/mL) or MDH (20 μM) to visualize LDs in imaging media prior to imaging. Confocal z-stacks were taken to sample total number of LDs throughout cells. To quantify total LD content, individual cells were randomly selected, and maximal intensity projections (MIPs) were created from z-stacks. Total LD area was segmented by intensity thresholding of BODIPY or MDH and measured using ‘Analyze Particles’ in Fiji. Relative LD contents were obtained by normalizing all data to the average of controls.

#### Mitochondrial FA import

WT and *OPA1* KO U2OS cells grown on 8-well chambered cover glass transduced with TOM20-Halo were treated with 20 µM of PF-04620110 (DGAT1 inhibitor) and 20 µM PF-06424439 (DGAT2 inhibitor) simultaneously or with 10 µM triacsin C in imaging media in the presence of 100 nM JF646 for 30 min to blunt LD biogenesis or mitochondrial FA trafficking, respectively. Without changing media, treated cells were then incubated with 1 µM TopFluor-OA, 1 µM BODIPY-linoleic acid, or 1 µM NBD-arachidonic acid for 10 min prior to imaging. Z-stack images from the bottom half of the cells were taken. To quantify mitochondrial FA import, MIPs from three continuous images from the bottom were first generated in Fiji.

Areas at the cell periphery were cropped (examples in Fig. S1H) to avoid out-of-focus signals in the peri-nuclear regions. Mitochondria masks were created by thresholding the TOM20-Halo channel and mean FA intensity within masks were measured.

*Relative enrichment of OPA1 on LD and in mitochondria.* To image OPA1 isoforms and mutants, cells grown in complete basal media on imaging cover glasses for 24 h were transduced with OPA1-expressing lentivirus. One day later, transduced cells were incubated with fresh media containing indicated FAs for another 20 h. Prior to imaging, cells were stained with 200 nM MitoTracker DeepRed for 3–5 min followed by media change to imaging media containing an LD stain (BODIPY or MDH). To quantify relative protein enrichment on LDs, confocal z-stack images of OPA1 isoforms and mutants as well as exon 4 and mutants were acquired with the Nikon confocal microscope together with LD staining and/or MitoTracker. MIPs of the LD and mitochondria channels were used to create LD and mitochondria masks via thresholding, which was then applied to the OPA1 channel using the region-of-interest manager for LD mean intensity (mean_LD_) and mitochondria mean intensity (mean_mito_) measurement. Mean intensity of the whole cell (mean_cell_) or of regions devoid of LDs (mean_cyt_) were also quantified for expression level normalization.

#### Relative enrichment of ATGL and CGI-58 on LDs

For ATGL imaging, U2OS cells were seeded onto imaging cover glass one day prior to transduction with ATGL S47A-Halo lentivirus at an MOI of 10–40. Cells were treated with OA 24 h after transduction and imaged with the Nikon confocal microscope 40 h later. For CGI-58 imaging, U2OS cells were transfected with 50 ng mApple-CGI-58 plasmid, treated overnight with 100 µM OA, and imaged the following day. Quantification of the relative enrichment of ATGL and CGI-58 on LDs were performed similarly to the analysis of OPA1 described above.

#### Inducible production of OPA1

*OPA1* KO U2OS cells were plated in 8-well chambered coverglass and transduced with TRE-OPA1-YFP-E3G or TRE-OPA1^iso2^-YFP-E3G lentivirus at an MOI of 100. After 48 h, transduced cells were incubated with 100 µM OA for 8 h to induce LD formation, followed by staining with 200 nM MitoTracker Red CMXRos for 5–10 min. Time-lapse imaging was conducted in 30-min intervals for 14 h in imaging media containing BODIPY-665/676 to stain LDs and 4 µg/mL doxycycline to induce OPA1 production. For the experiment looking at the absence or presence (±) of pyruvate, *OPA1* KO U2OS cells were transduced with TRE-OPA1-YFP-E3G for 24 h, followed by a change of media containing 100 µM OA and 0–1 mM pyruvate. Approximately 20 h later, 4 µg/mL doxycycline was added for 4–5 h with indicated levels of pyruvate to induce OPA1-YFP expression prior to imaging. OPA1-YFP or OPA1^iso2^-YFP signal within individual cells were categorized into: (i) mitochondria, (ii) LDs, or (iii) mitochondria and LDs, based on their subcellular distribution.

#### Interaction between mitochondria and LD

U2OS cells and U2OS were transduced with lentiviral OPA1-YFP at an MOI=100 and incubated with 100 μM OA overnight as described above. To quantify mitochondria and LD colocalization, MIPs of mitochondria and LDs channels were created in Fiji, and their colocalization was analyzed using the JACoP plugin^107^. To analyze the relative contact sites between mitochondria and LDs, WT and *OPA1* KO U2OS cells were transduced with lentivirus expressing the Mito-LD FABCON reporter at a MOI of 300. Cells were imaged and analyzed as described previously^60^.

#### Mitochondrial morphology analysis

U2OS WT, *OPA1* KO, and *OPA1* KO cells reconstituted with OPA1 mutants across indicated conditions were stained with 200 nM MitoTracker Deep Red for 3 min. Cells were washed in warm McCoy’s 5A medium three times before Z-stack images were acquired. To quantify mitochondrial morphological features, MIPs were generated from three continuous bottom images and analyzed using Fiji’s ‘Mitochondria-Analyzer’ plug-in^66^. The 2D analysis mode was used with a block size of 1.350 microns and C-value of 5.

#### Correlative confocal-scanning electron microscopy

Four thousand U2OS cells in 300 µL McCoy’s 5A medium were plated onto the gridded cover glass area within MatTek dishes (Cat# P35G-1.5-14-C-GRID, MatTek). Twenty-four hours after plating, cells were transduced with lenti-OPA1-mhYFP at an MOI of 50. Twenty-four hours after transduction, the media was changed to include 100 µM OA and left to incubate overnight. Alternatively, for the exon 4 experiment, cells were transfected with 150 ng exon4-hfYFP plasmid 48 h after plating in fresh media with 100 µM OA overnight. Transduced or transfected cells were sequentially stained with 400 nM MitoTracker Red CMXRos and 20 µM Hoechst at 37°C for 5 min, followed by staining with 500 ng/mL BODIPY 665 at 37°C for 30 min. After staining, cells were fixed with 2.5% glutaraldehyde and 2% PFA fixation in 0.1 M sucrose prepared in 0.1 M Sorenson’s phosphate buffer (PB) at room temperature for 30 min. Fixed cells were rinsed before imaging in PBS to acquire z-stack fluorescence images and locate cells of interest.

After confocal imaging, cells were further fixed in 2.5% glutaraldehyde, 2% PFA, and 0.1 M sucrose in PB for 3 days at 4°C. The fixed cells were post-fixed in a mixture of 0.5% OsO_4_ + 0.5% K_4_Fe(CN)_6_ in 0.1 M PB for 10 min on Lab Armor beads pre-chilled to 4°C in a Styrofoam cooler for lipid fixation and contrasting. The post-fixed cells were further contrasted with 1% uranyl acetate in H_2_O overnight at 4°C. Cells were dehydrated with an ascending ethanol series (10%, 30%, 50%, 70%, 80%, 90%, 95%, and 100%), with each step carried out for 5 min on cold beads, before the cells were embedded in Spurr’s resin (Cat#14300, Electron Microscopy Sciences). The resin-embedded cells were thermally polymerized for 24 h at 70°C. After thermal polymerization, all cells, including the target cells, were transferred to the polymerized resin side of the MatTek dish without any physical damage once the glass cover slip was removed.

Following transfer, the target cells were serially sectioned to a dimension of 300 µm x 300 µm x 100 nm on a Leica ARTOS 3D ultramicrotome using a DiATOME Histo Jumbo diamond knife. Approximately 40 serial sections with the desired dimension were collected on a silicon chip (5 mm x 7 mm) and dried for 1 h at 45°C. The serial sections were coated with 6 nm carbon in a Leica EM ACE600 High Vacuum Coater before scanning electron microscope (SEM)-based array tomography imaging. The target cells were imaged in a Zeiss GeminiSEM 460 using a Sense backscattered electron detector with 1.5 kV high tension and 80 pA current. Zeiss Atlas 5 Array Tomography software assisted in the automatic selection of the target cells throughout the serial sections and performed high-resolution imaging at 2-nm pixel resolution in a 30 µm x 30 µm of field of view. The obtained images from serial sections were aligned using the lipid droplets as internal fiducials and exported as tagged image file format (TIFF) files by Atlas 5 Array Tomography to co-register to the obtained confocal microscopic images.

### Prediction of OPA1 MTS strength

The MTS of OPA1 was tested for its relative strength by calculating its amphiphilicity as described^58^. Amphiphilicity was calculated by defining the net charge of the OPA1 MTS at pH 7.4 (calculated at ProtPI, https://www.protpi.ch/Calculator/ProteinTool) and its maximal hydrophobic moment (calculated at EMBOSS, https://www.bioinformatics.nl/cgi-bin/emboss/hmoment) and multiplying these two values. This amphiphilicity value was compared with experimentally determined parameters defining MTS strength for 16 independent constructs, as reported^58, 59^. The OPA1 MTS was defined based on a previous study^28^ and was additionally defined through the MitoFates database (https://mitf.cbrc.pj.aist.go.jp/MitoFates/cgi-bin/top.cgi).

### GTEx data mining

All GTEx (V8) RNA-seq data were remapped to human genome reference GRCh38 using Gencode reference annotation with STAR (v2.7.9a)^108^ in a splice-aware manner. The junction read counts for *OPA1* Exon3-Exon4, Exon 4-Exon5 and Exon3-Exon5 were used to compute the percent of inclusion and PSE of exon 4.

### SJLIFE data analysis

The St. Jude Lifetime cohort participants return for a clinical visit that includes sample collection, germline genotyping and laboratory testing^71^.

This analysis focused on replication of association signals for a single germline variant, chr3:193617202:G>A (hg38), in five prespecified lipid related traits in SJLIFE: body fat percentage, blood TAGs, total cholesterol, low-density lipoprotein (LDL) cholesterol, and HDL cholesterol. Whole-exome sequencing data from 4,328 SJLIFE participants with genotype data were used as the starting dataset before trait-specific phenotype missingness was applied. The cohort was 52.1% male, and the mean age at sample collection was 27.99 years (SD 11.39; median 27.13).

For each of the five traits, we first evaluated whether analysis should be performed on the raw phenotype scale or after inverse-rank normal transformation (IRNT), based on the distribution of covariate-adjusted residuals. Raw values were retained only when absolute residual skewness ≤ 0.5 and absolute excess kurtosis value ≤ 1; otherwise IRNT was used. Under this rule, body fat percentage was analyzed on the raw scale, while bloodTAGs, total cholesterol, LDL cholesterol, and HDL cholesterol were analyzed on the IRNT scale.

Genotypic associations were evaluated using ordinary least-squares regression in Python (statsmodels), with separate indicators for G/A and A/A and G/G as the reference genotype. Models were adjusted for age at last contact, age at diagnosis, the recorded Gender variable, receipt of chemotherapy and radiation, and the first five genetic ancestry principal components (PC1–PC5). G/A-vs-G/G and A/A-vs-G/G contrasts were evaluated using two-sided *t*-tests, and the overall genotype association was evaluated using a joint two-degrees-of-freedom Wald χ² test.

### Statistical analysis

Raw data points (semi-transparent symbols) and mean ± standard deviation from independent experiments (bold symbols and lines) were plotted together to simultaneously convey cell variability and experimental reproducibility. Mean values from independent experiments were statistically analyzed in GraphPad Prism (version 11.0.0, GraphPad Software, Inc., San Diego, CA, USA) using unpaired or paired *t*-test or one-way ANOVA. Graphs and figures were generated using GraphPad Prism and Adobe Illustrator (version 29.8.5, San Jose, CA, USA).

## Acknowledgements

We would like to thank Caleb Lindow for generating some of the OPA1 truncation plasmids, Drs. Mondira Kundu and Kirsten K. Ness for consultation on the St. Jude LIFE data, Dr. Danny M. D’Amore for scientific editing, Drs. Ashley Jones and Meghen McGehee for harvesting mouse livers, Dr. Lauren Brakefield-Laird for technical consultations, Justine Zhao for technical assistance, and Gunda Johnson, Marquetta Nebo, Jennifer Maglisco, and Monique Payton for administrative and operational assistance. This research included experiments conducted by St. Jude Comprehensive Cancer Center Shared Resources which are supported in part by the National Cancer Institute grant P30 CA021765. This work was supported by ALSAC (to C-L.C.) and the National Institutes of Health (1DP2GM150192 to C-L.C. and R35GM151130 to N.M.N.).

## Author contributions

X.L. and C-L.C. conceived the project, performed image acquisition, image analysis, and structure predictions. X.L., M.R.H, and R.B. performed the LD fractionation and western blotting. D.V. performed FAO assays and analyses, which were supervised by J.T.O. J.K. generated and validated the U2OS *OPA1* KO cells, which was supervised by S.P-M. M.H. and X.L. performed the RT-PCR. W.J.C. performed the CLEM imaging with supervision by C.G.R. R.E.T. supervised the generation of all the lentiviral vectors. G.W. performed the exon 4 splicing analysis from the GTEx database. C.L. and Y.S. performed the patient data analysis from the St. Jude LIFE cohort database. N.M.N. performed the mitochondrial import analysis for OPA1. X.L. and C-L.C wrote the manuscript with input from all authors. C-L.C supervised the project.

---

**Figure S1.**
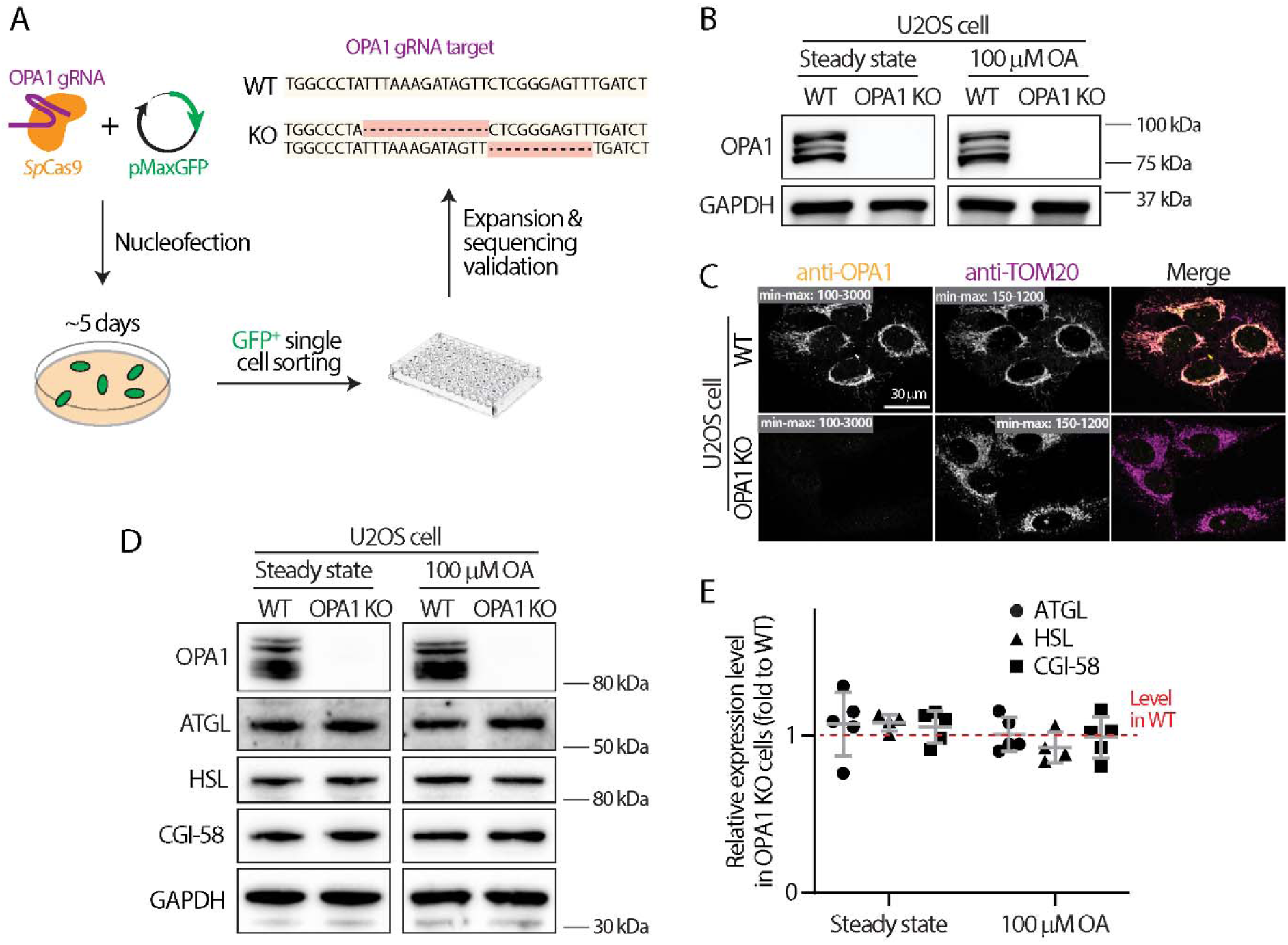
Generation and validation of *OPA1* knockout and knockdown cells. **(A)** Diagram illustrating the generation of *OPA1* knockout (KO) U2OS cells via CRISPR-genome editing. Sequencing data validating *OPA1* KO is shown in the top right, and deleted *OPA1* genome sequences are represented as dashed lines highlighted in red. **(B)** Western blot analysis of OPA1 and GAPDH in wildtype (WT) and *OPA1* KO U2OS cells ± overnight 100-µM oleic acid (OA) treatment. **(C)** Immunostaining of endogenous OPA1 and TOM20 in WT and *OPA1* KO U2OS cells detected by confocal microscopy. Maximal intensity projected confocal images from whole cells with min-max intensity range (gray boxes) are shown. **(D)** Western blot analysis of ATGL, HSL, CGI-58, and GAPDH in WT and *OPA1* KO U2OS cells ± overnight 100-µM OA treatment. **(E)** Quantification of GAPDH-normalized ATGL, HSL, and CGI-58 levels in *OPA1* KO U2OS cells in relative to WT cells at steady state or with overnight 100-µM OA treatment. Mean ± standard deviation from 4–5 independent experiments are shown. The differences are non-significant for all proteins, as assessed by unpaired *t*-test.

**Figure S2.**
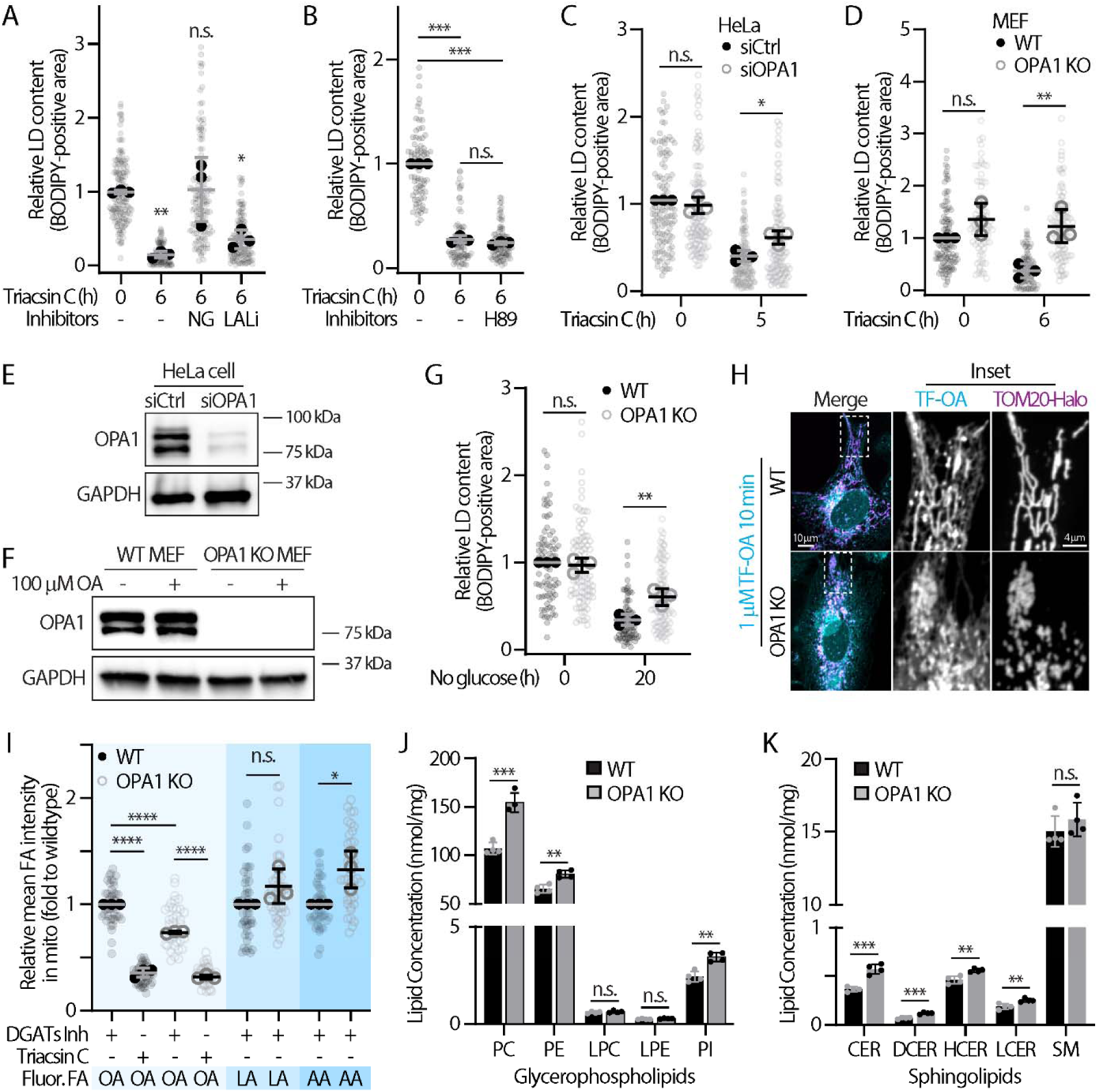
Characterization of the breakdown and biogenesis of lipid droplets and phospholipids in wildtype and *OPA1*-silenced cells. **(A)** Relative lipid droplet (LD) content (BODIPY 493/503-positive area) in widetype (WT) U2OS cells treated with 100 µM oleic acid (OA) overnight followed by 10 µM triacsin C incubation in the presence of 10 µM NG497 (NG) or 5 µM Lalistat 2 (LALi) for 6 h. Mean ± standard deviation from three independent experiments are shown (n = 129–148 cells). For statistics in panels A–E, n.s., no significance, ***P ≤ 0.001, **P ≤ 0.01, *P ≤ 0.05, as assessed by one-way ANOVA. **(B)** Relative LD content in U2OS cells treated with 100 µM OA overnight followed by 6-h co-incubation of 10 µM triacsin C and 10 µM H89. **(C)** Relative LD content in HeLa cells transfected with scramble siRNA (siCtrl) or *OPA1* siRNA (siOPA1) treated with 100 µM OA overnight before 6-h incubation with 10 µM triacsin C. Mean ± standard deviation from three independent experiments are shown (total of 107–122 cells). **(D)** Relative LD content in WT and *OPA1* knockout (KO) mouse embryonic fibroblasts (MEFs) treated with 100 µM OA overnight before 6-h incubation with 10 µM triacsin C. Mean ± standard deviation from three independent experiments are shown (n = 83–105 cells). **(E)** Western blot analysis of OPA1 and GAPDH in HeLa cells transfected with scramble siRNA (siCtrl) or OPA1 siRNA. **(F)** Western blot analysis of OPA1 and GAPDH in WT and *OPA1* KO MEFs ± overnight 100-µM OA treatment. **(G)** Relative LD content in WT and *OPA1* KO U2OS cells treated with 100 µM OA overnight followed by 20-h incubation in glucose-free Dulbecco’s Modified Eagle Medium. Mean ± standard deviation from three independent experiments are shown (n = 79–87 cells). **(H)** Subcellular localization of TopFluor (TF)-OA and TOM20-Halo (stained with JF646-HTL) detected via confocal microscopy in WT and *OPA1* KO U2OS cells treated with 20 µM DGAT1 and DGAT2 inhibitors. Maximal intensity projected confocal images from three axial slices (∼0.6 µm total thickness) are shown. **(I)** Quantification of the intensity of peripheral mitochondria (mito) of TF-OA as described in (F) and in cells pretreated with 10 µM triacsin C, 1 µM BODIPY-linoleic acid (LA), or 1 µM NBD-arachidonic acid (AA). Mean ± standard deviation from three independent experiments are shown (n = 38–50 cells). n.s., no significance, ****P ≤ 0.0001, *P ≤ 0.05, as assessed by one-way ANOVA. **(J and K)** Levels of (J) glycerophospholipids and (K) sphingolipids in steady-state WT and *OPA1* KO U2OS cells determined using liquid chromatography–mass spectrometry. Mean ± standard deviation from four replicates are shown. n.s., no significance, ***P ≤ 0.001, **P ≤ 0.01, as assessed by unpaired *t*-test. Abbreviations: PC, phosphatidylcholine; PE, phosphatidylethanolamine; LPC, lysophosphatidylcholine; LPE, lysophosphatidylethanolamine; PI, phosphatidylinositol; CER, ceramide; DCER, dihydroceramide; HCER, hexosylceramide; LCER, lactosylceramide; SM, sphingomyelin.

**Figure S3.**
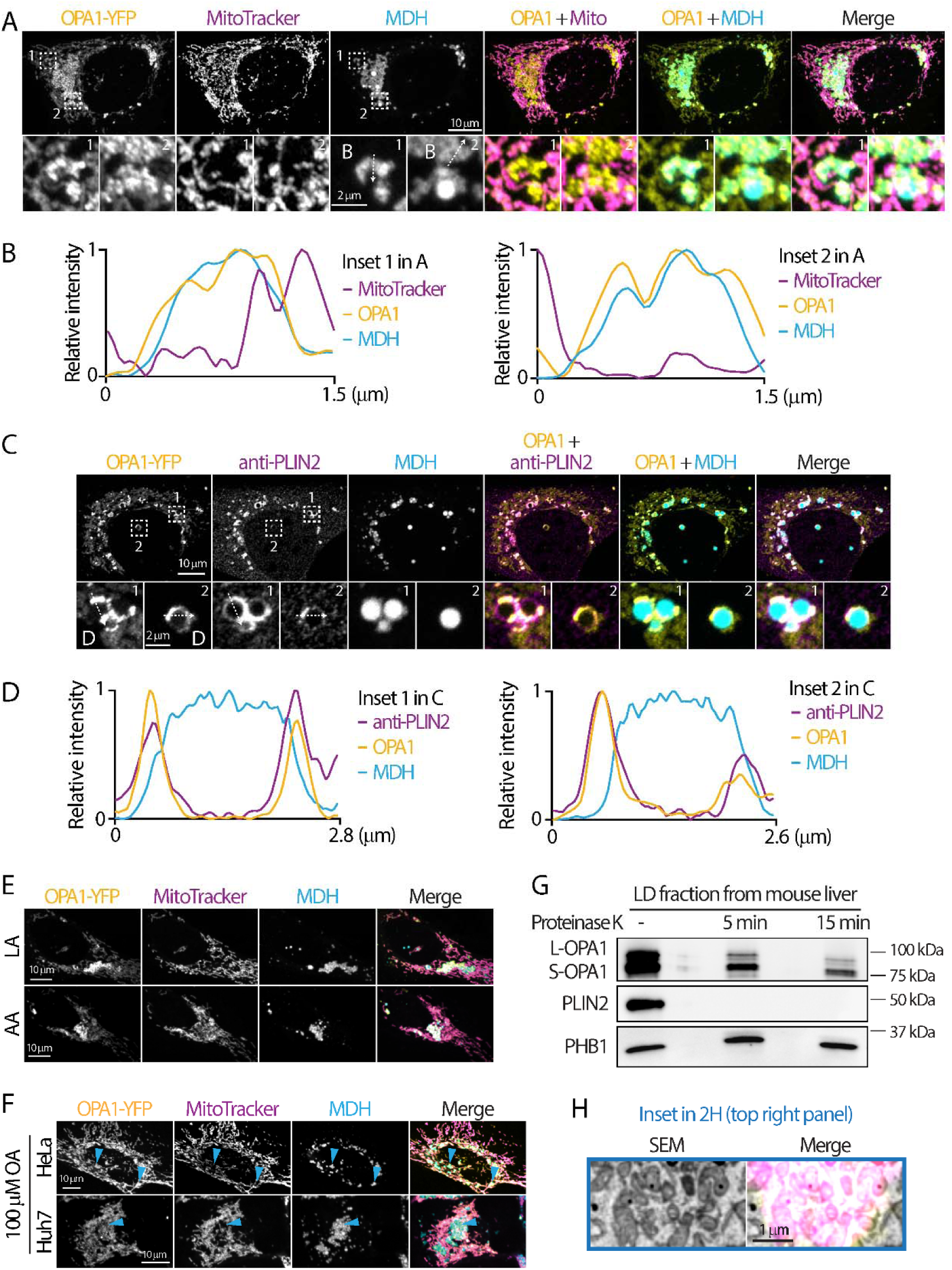
Localization of OPA1 on lipid droplets. **(A)** Subcellular localization of OPA1-YFP (lentiviral), mitochondria (MitoTracker Deep Red), and lipid droplets (LDs; MDH) by Airyscan microscopy in fixed U2OS cells treated with 100 µM oleic acid (OA) overnight. Maximal intensity projected (MIP) of Airyscan images from 17 axial slices (∼2.4 µm total thickness) are shown. **(B)** Relative intensity profiles of OPA1, MitoTracker, and MDH measured along the white-dashed arrow from inset 1 and 2 in (A). **(C)** Subcellular localization of OPA1-YFP (lentiviral), mitochondria (MitoTracker Deep Red), and an endogenous LD protein (anti-PLIN2) by Airyscan microscopy in fixed U2OS cells treated with 100 µM OA overnight. MIP of Airyscan images from 18 axial slices (∼2.2 µm total thickness) are shown. **(D)** Relative intensity profiles of OPA1, MitoTracker, and PLIN2 measured along the white-dashed arrow from inset 1 and 2 in (C). **(E)** Localization of OPA1-YFP (lentiviral), mitochondria (MitoTracker Deep Red), and LDs (MDH) in U2OS cells treated with 100 µM linoleic acid (LA) or 100 µM arachidonic acid (AA) overnight. MIP confocal images from four axial slices (∼1 µm total thickness) are shown. **(F)** Localization of OPA1-YFP, mitochondria (MitoTracker Deep Red), and LDs (MDH) in HeLa and Huh7 cells treated with 100 µM OA overnight, detected with confocal microscopy. MIP confocal images from four axial slices (∼1 µm total thickness) are shown. Cyan arrowheads indicate area with LDs. **(G)** Western blot analysis of OPA1, PLIN2, and PHB1 in isolated LD fractions from mouse livers before and after treatment with 10 µg/ml of proteinase K treatment for 5 or 15 min. **(H)** Scanning electron microscopy (SEM) and SEM merged with OPA1 fluorescence images from blue inset in Fig. 2H.

**Figure S4.**
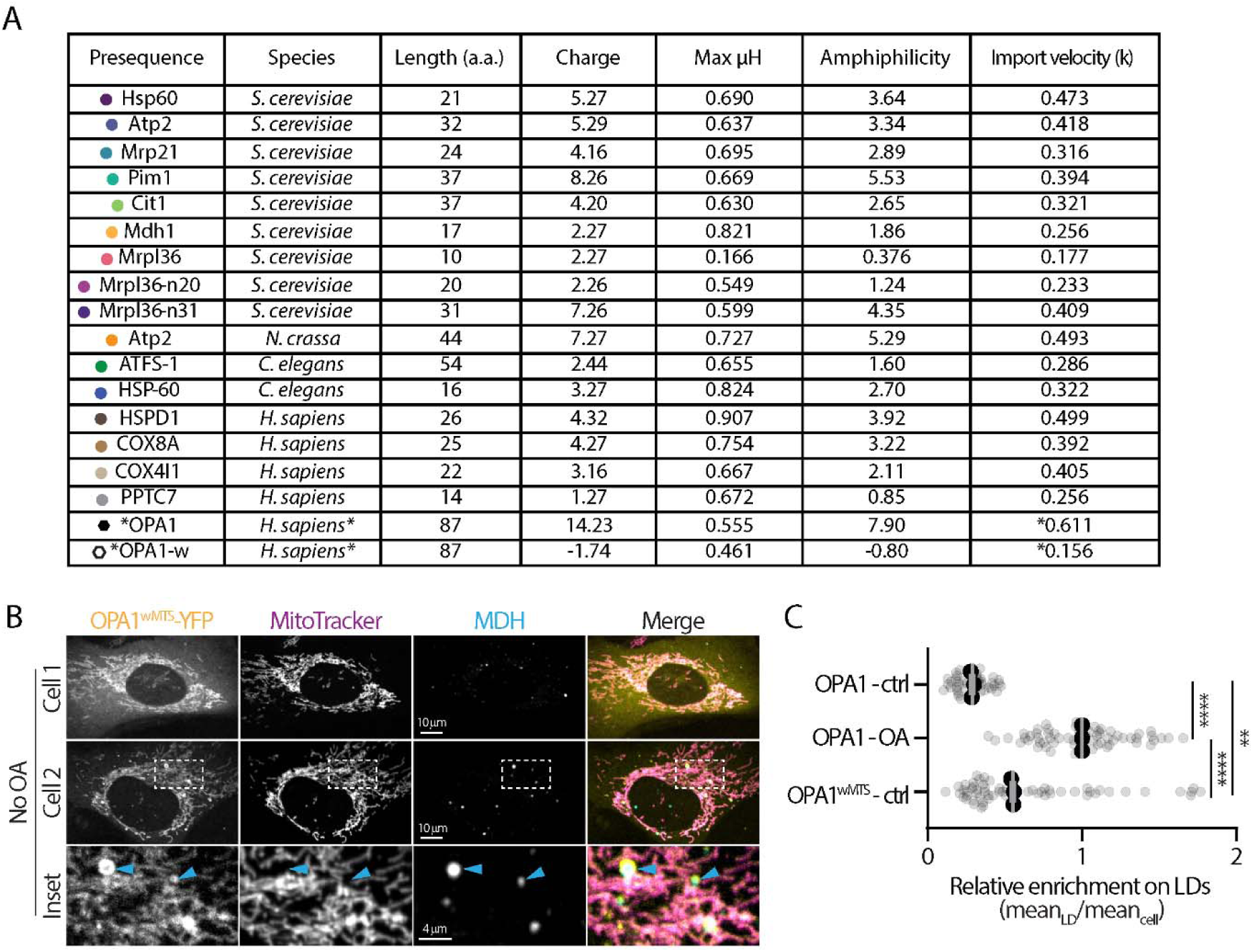
Prediction of OPA1 import velocity and localization of OPA1^wMTS^. **(A)** Properties and experimental determined import velocity (k) of 16 mitochondrial targeting sequences (MTSs) and predicted (indicated by asterisk) import velocity of the MTSs of OPA1 and OPA1^wMTS^. Abbreviation: Max μH, maximal helical hydrophobic moment. **(B)** Localization of OPA1^wMTS^-YFP, mitochondria (Mitotracker Deep Red), and lipid droplets (LDs; MDH) in U2OS cells without oleic acid (OA) treatment. **(C)** Relative enrichment of OPA1^wMTS^ from (B) and OPA1 on LDs. Means ± standard deviation from three independent experiments are shown (total of 45–63 cells). ****P ≤ 0.0001, **P ≤ 0.01, as assessed by one-way ANOVA.

**Figure S5.**
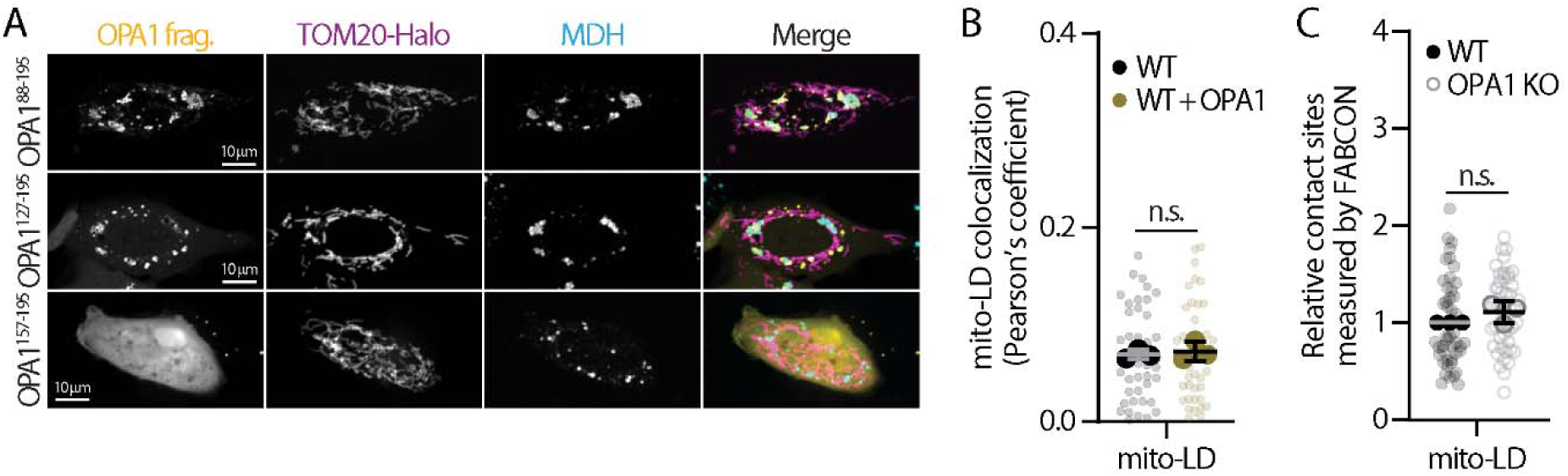
Localization of OPA1 truncations and quantification of mitochondria–lipid droplet contact sites. **(A)** Localization of truncated YFP-tagged OPA1 fragments (frag.), mitochondria (labeled with TOM20-Halo stained with JF646-HTL), and lipid droplets (LDs; MDH) in U2OS cells treated with 100 µM oleic acid overnight and detected by confocal microscopy. Maximal intensity projected confocal images from four axial slices (∼1 µm total thickness) are shown. **(B)** Quantification of colocalization between mitochondria and LDs in wildtype (WT) and OPA1-YFP–overexpressing U2OS cells. Means ± standard deviation from three independent experiments are shown (total of 41–42 cells). n.s., no significance, as assessed by paired *t*-test. **(C)** Relative mitochondria-LD contact sites measured by FABCON reporter in WT and *OPA1* knockout (KO) U2OS cells. Means ± standard deviation from three independent experiments are shown (total of 47–49 cells). n.s., no significance, as assessed by paired *t*-test.

**Figure S6.**
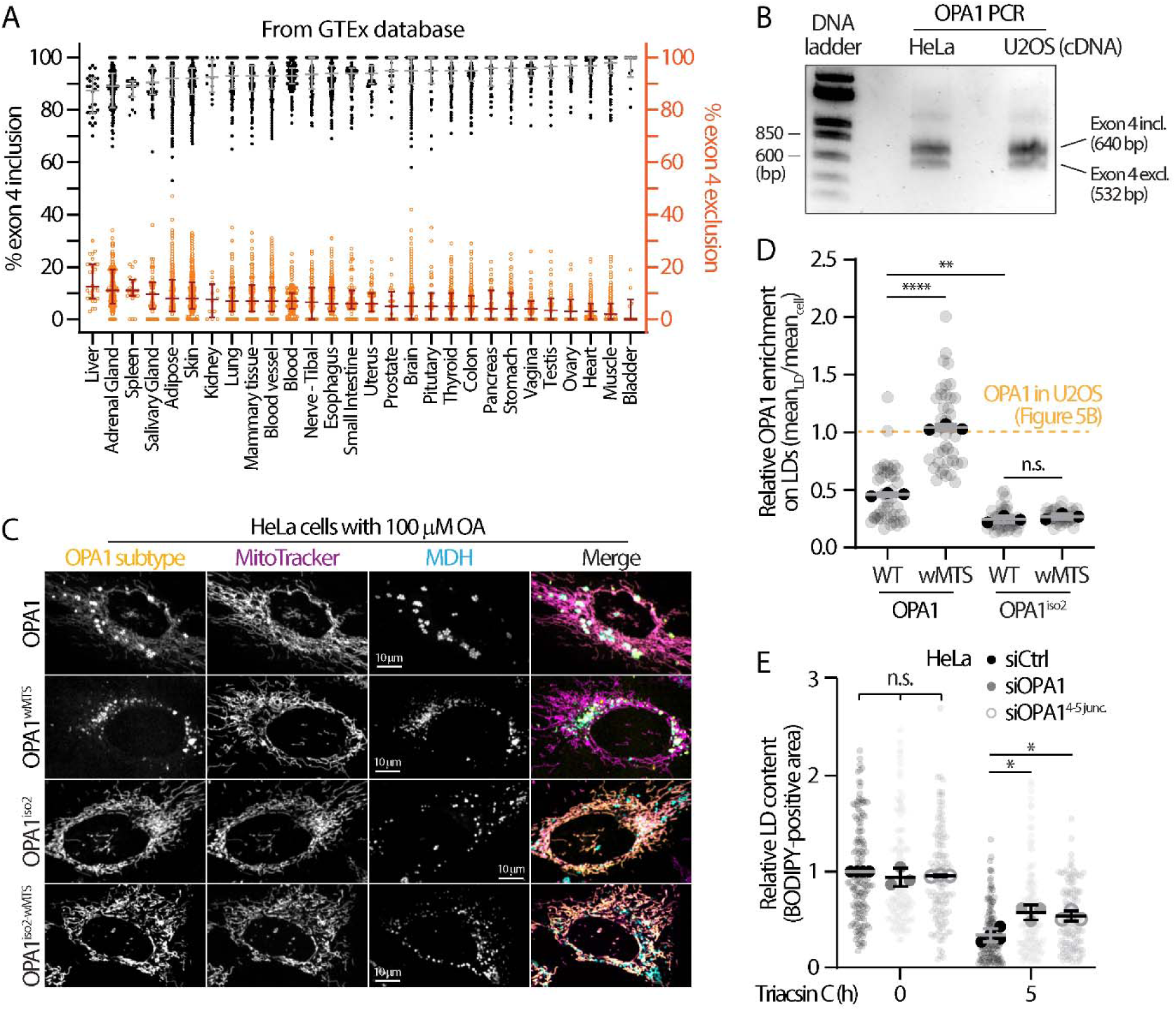
OPA1^iso2^ mRNA expression, protein localization, and effect on fatty acid release. **(A)** Percentage of OPA1 exon 4 inclusion and exclusion, representing OPA1 isoform 1 and isoform 2, respectively, across various human tissues analyzed from the Genotype-Tissue Expression (GTEx) database. Raw data and median values with quartile ranges are shown. **(B)** Profiling of OPA1 exon 4 inclusion (incl.) and exclusion (excl.) from cDNA of HeLa and U2OS cells. **(C)** Localization of OPA1-YFP, OPA1^wMTS^-YFP, OPA1^iso2^-YFP, and OPA1^iso2-wMTS^-YFP, mitochondria (MitoTracker Deep Red), and lipid droplets (LDs; MDH) in HeLa cells treated with 100 µM oleic acid (OA) overnight and detected by confocal microscopy. Maximal intensity projected confocal images from four axial slices (∼1 µm in total thickness) are shown. **(D)** Relative enrichment of OPA1 and variants on LDs as described in (C). Means ± standard deviation from three or four independent experiments are shown (total of 40–42 cells). Yellow dashed line indicates the relative enrichment of OPA1 on LDs in U2OS cells as described in Fig. 5B. n.s., no significance, ****P ≤ 0.0001, **P ≤ 0.01, as assessed by one-way ANOVA. **(E)** BODIPY-positive LD content in siCtrl-, siOPA1-, and siOPA1^4-5^ ^junc^-transfected HeLa cells treated with 100 µM OA overnight followed by incubation with 10 µM triacsin C for 5 h. Means ± standard deviation from 3–4 independent experiments are shown (total of 117–156 cells). n.s., no significance, *P ≤ 0.05, as assessed by one-way ANOVA.

**Figure S7.**
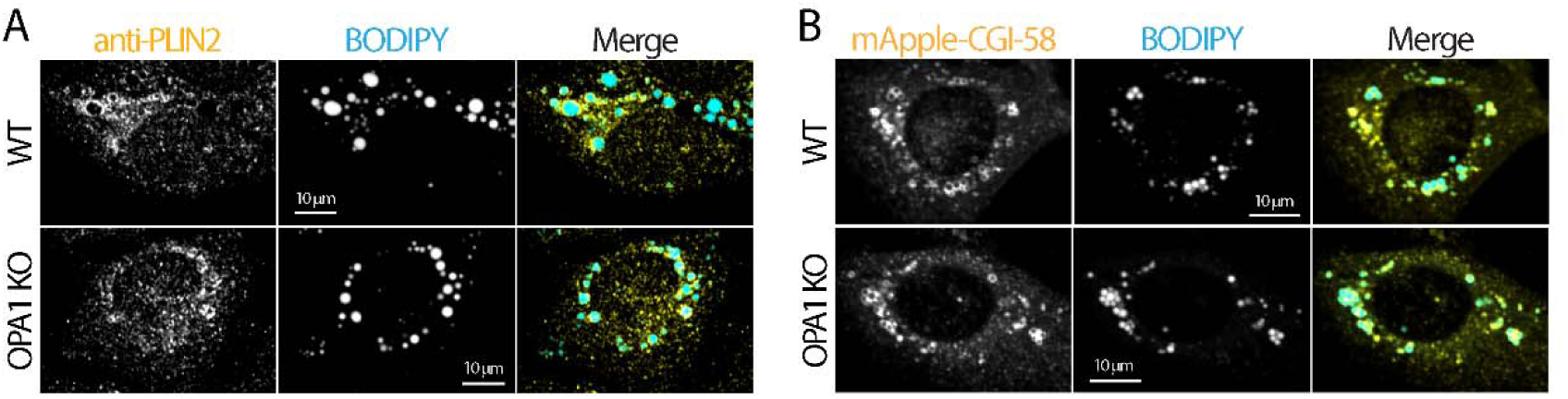
Lipid droplet protein distributions in wildtype and *OPA1* knockout U2OS cells. **(A)** Localization of endogenous PLIN2 on lipid droplets (LDs) labeled with BODIPY-493/503 in wildtype (WT) and *OPA1* knockout (KO) U2OS cells treated with 100 µM oleic acid (OA) overnight. Maximal intensity projected (MIP) confocal images from four axial slices (∼1 µm total thickness) are shown. **(B)** Localization of mApple-CGI-58 and LDs labeled with BODIPY-493/503 in WT and *OPA1* KO U2OS cells treated with 100 µM OA overnight. MIP confocal images from four axial slices (∼1 µm total thickness) are shown.

**Figure S8.**
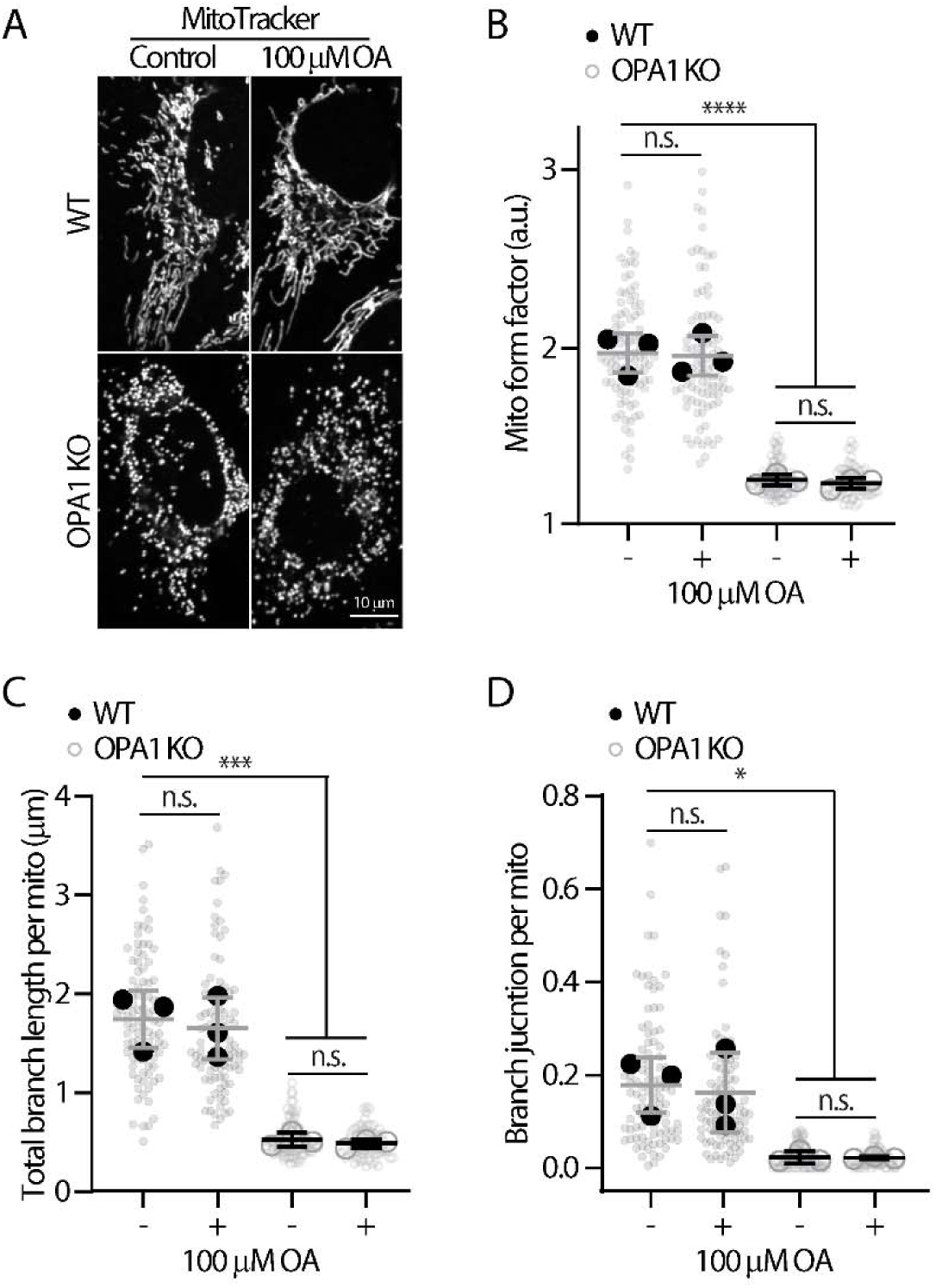
Mitochondrial characteristics in wildtype and *OPA1* knockout U2OS cells. **(A)** Representative images of mitochondria stained with MitoTracker DeepRed in wildtype (WT) and *OPA1* knockout (KO) U2OS cells ± overnight 100 µM oleic acid (OA) treatment. MIP confocal images from four axial slices (∼1 µm total thickness) are shown. **(B–D)** 2D ‘mitochondria analyzer’ analysis of mitochondrial morphology and connectivity measuring (B) form factor, (C) total branch length, and (D) branch junction in MitoTracker DeepRed–stained WT and *OPA1* KO U2OS cells ± overnight 100 µM oleic acid treatment. Means ± standard deviation from three independent experiments are shown (total of 74–100 cells). n.s., no significance, ****P ≤ 0.0001, ***P ≤ 0.001, *P ≤ 0.05, as assessed by one-way ANOVA.

