## Supplementary material for "Alternative organelle targeting of OPA1 mediates fatty acid release from lipid droplets": Table S1

| gnomAD ID | rsIDs | Reference | Alternate | HGVS Consequence | Protein Consequence | Transcript Consequence | VEP Annotation | Allele Frequency (high to low) |
|---|---|---|---|---|---|---|---|---|
| 3-193617202-G-A | rs7624750 | G | A | p.Ser158Asn | p.Ser158Asn | c.473G>A | missense_variant | 0.47097541 |
| 3-193617259-G-T | rs150279202 | G | T | p.Ser177Ile | p.Ser177Ile | c.530G>T | missense_variant | 0.000106719 |
| 3-193617246-G-A | rs778997114 | G | A | p.Val173Ile | p.Val173Ile | c.517G>A | missense_variant | 6.50941E-05 |
| 3-193617234-TTTG-T | rs760263649 | TTTG | T | p.Phe169_Asp170delins[illegible] | p.Phe169_Asp170delins[illegible] | c.506_508del | inframe_deletion | 3.59527E-05 |
| 3-193617229-C-T | rs754177232 | C | T | p.Pro167Leu | p.Pro167Leu | c.500C>T | missense_variant | 1.79778E-05 |
| 3-193617184-T-A | rs2108877878 | T | A | p.Ile152Asn | p.Ile152Asn | c.455T>A | missense_variant | 1.19458E-05 |
| 3-193617264-T-A | | T | A | p.Leu179Met | p.Leu179Met | c.535T>A | missense_variant | 5.58791E-06 |
| 3-193617273-T-C | rs769335936 | T | C | p.Phe182Leu | p.Phe182Leu | c.544T>C | missense_variant | 4.9951E-06 |
| 3-193617193-C-T | rs751318725 | C | T | p.Ala155Val | p.Ala155Val | c.464C>T | missense_variant | 3.12227E-06 |
| 3-193617279-A-G | | A | G | p.Thr184Ala | p.Thr184Ala | c.550A>G | missense_variant | 1.88651E-06 |
| 3-193617186-A-G | rs762882041 | A | G | p.Arg153Gly | p.Arg153Gly | c.457A>G | missense_variant | 1.88238E-06 |
| 3-193617260-C-A | rs727504057 | C | A | p.Ser177Arg | p.Ser177Arg | c.531C>A | missense_variant | 1.86165E-06 |
| 3-193617252-A-C | rs1729174080 | A | C | p.Ser175Arg | p.Ser175Arg | c.523A>C | missense_variant | 1.86017E-06 |
| 3-193617235-T-C | rs1386915863 | T | C | p.Phe169Ser | p.Phe169Ser | c.506T>C | missense_variant | 1.8598E-06 |
| 3-193617237-G-A | | G | A | p.Asp170Asn | p.Asp170Asn | c.508G>A | missense_variant | 1.85974E-06 |
| 3-193617221-G-C | rs1380881656 | G | C | p.Lys164Asn | p.Lys164Asn | c.492G>C | missense_variant | 1.85966E-06 |
| 3-193617233-C-G | rs1262531072 | C | G | p.Asp168Glu | p.Asp168Glu | c.504C>G | missense_variant | 1.85957E-06 |
| 3-193617187-G-A | rs766257852 | G | A | p.Arg153Lys | p.Arg153Lys | c.458G>A | missense_variant | 1.25197E-06 |
| 3-193617271-A-G | rs138884045 | A | G | p.Asp181Gly | p.Asp181Gly | c.542A>G | missense_variant | 1.24537E-06 |
| 3-193617269-G-T | rs747354358 | G | T | p.Lys180Asn | p.Lys180Asn | c.540G>T | missense_variant | 1.24328E-06 |
| 3-193617201-A-G | | A | G | p.Ser158Gly | p.Ser158Gly | c.472A>G | missense_variant | 1.24267E-06 |
| 3-193617228-C-T | rs1729170377 | C | T | p.Pro167Ser | p.Pro167Ser | c.499C>T | missense_variant | 1.23977E-06 |
| 3-193617229-C-G | rs754177232 | C | G | p.Pro167Arg | p.Pro167Arg | c.500C>G | missense_variant | 1.23976E-06 |
| 3-193617224-A-C | | A | C | p.Leu165Phe | p.Leu165Phe | c.495A>C | missense_variant | 1.23968E-06 |
| 3-193617181-A-G | rs1329178588 | A | G | p.Lys151Arg | p.Lys151Arg | c.452A>G | missense_variant | 6.33E-07 |
| 3-193617183-A-T | | A | T | p.Ile152Phe | p.Ile152Phe | c.454A>T | missense_variant | 6.29E-07 |
| 3-193617280-C-T | | C | T | p.Thr184Ile | p.Thr184Ile | c.551C>T | missense_variant | 6.29E-07 |
| 3-193617277-T-C | rs772737325 | T | C | p.Phe183Ser | p.Phe183Ser | c.548T>C | missense_variant | 6.27E-07 |
| 3-193617274-T-G | | T | G | p.Phe182Cys | p.Phe182Cys | c.545T>G | missense_variant | 6.24E-07 |
| 3-193617196-T-G | | T | G | p.Leu156Arg | p.Leu156Arg | c.467T>G | missense_variant | 6.22E-07 |
| 3-193617198-C-T | | C | T | p.Pro157Ser | p.Pro157Ser | c.469C>T | missense_variant | 6.22E-07 |
| 3-193617270-G-C | rs780818428 | G | C | p.Asp181His | p.Asp181His | c.541G>C | missense_variant | 6.22E-07 |
| 3-193617268-A-C | rs780451120 | A | C | p.Lys180Thr | p.Lys180Thr | c.539A>C | missense_variant | 6.21E-07 |
| 3-193617266-G-T | | G | T | p.Leu179Phe | p.Leu179Phe | c.537G>T | missense_variant | 6.21E-07 |
| 3-193617263-A-C | | A | C | p.Leu178Phe | p.Leu178Phe | c.534A>C | missense_variant | 6.21E-07 |
| 3-193617207-G-A | | G | A | p.Glu160Lys | p.Glu160Lys | c.478G>A | missense_variant | 6.20E-07 |
| 3-193617254-C-G | | C | G | p.Ser175Arg | p.Ser175Arg | c.525C>G | missense_variant | 6.20E-07 |
| 3-193617255-C-G | rs1729174512 | C | G | p.Leu176Val | p.Leu176Val | c.526C>G | missense_variant | 6.20E-07 |
| 3-193617216-G-A | | G | A | p.Val163Ile | p.Val163Ile | c.487G>A | missense_variant | 6.20E-07 |
| 3-193617244-T-C | | T | C | p.Ile172Thr | p.Ile172Thr | c.515T>C | missense_variant | 6.20E-07 |
| 3-193617235-T-G | | T | G | p.Phe169Cys | p.Phe169Cys | c.506T>G | missense_variant | 6.20E-07 |
| 3-193617238-A-T | | A | T | p.Asp170Val | p.Asp170Val | c.509A>T | missense_variant | 6.20E-07 |
| 3-193617231-G-C | | G | C | p.Asp168His | p.Asp168His | c.502G>C | missense_variant | 6.20E-07 |
