## Supplementary material for "Alternative organelle targeting of OPA1 mediates fatty acid release from lipid droplets": Table S2

| Trait | Genotype | N | Mean_native | SEM_native | adj_difference_ | p_inference | joint_2df_p |
|---|---|---|---|---|---|---|---|
| Body fat | G/G | 903 | 34.6297935 | 0.28783601 | | | 0.09333569 |
| Body fat | G/A | 1759 | 34.2530436 | 0.21319151 | -0.3998409 | 0.16857031 | 0.09333569 |
| Body fat | A/A | 843 | 35.1262583 | 0.30401969 | 0.20877811 | 0.53882218 | 0.09333569 |
| Blood TAG | G/G | 1104 | 126.396739 | 3.84793683 | | | 0.00334292 |
| Blood TAG | G/A | 2171 | 120.168125 | 2.5310439 | -7.1634078 | 0.01883196 | 0.00334292 |
| Blood TAG | A/A | 1052 | 118.38308 | 4.7099399 | -7.6425412 | 0.0008734 | 0.00334292 |
| Blood cholesterol | G/G | 1104 | 179.986413 | 1.19333292 | | | 0.03959379 |
| Blood cholesterol | G/A | 2171 | 176.777982 | 0.85276382 | -3.3279415 | 0.01400596 | 0.03959379 |
| Blood cholesterol | A/A | 1052 | 176.811787 | 1.22799382 | -2.9187839 | 0.05395327 | 0.03959379 |
| LDL | G/G | 1096 | 103.898723 | 1.0291504 | | | 0.14600618 |
| LDL | G/A | 2158 | 101.539388 | 0.71370728 | -2.486674 | 0.05199333 | 0.14600618 |
| LDL | A/A | 1043 | 102.206136 | 1.0807206 | -1.4189612 | 0.37518609 | 0.14600618 |
| HDL | G/G | 1104 | 51.673913 | 0.4995026 | | | 0.71336996 |
| HDL | G/A | 2171 | 51.7678489 | 0.34745108 | 0.27428708 | 0.42893753 | 0.71336996 |
| HDL | A/A | 1052 | 51.8602662 | 0.47349121 | 0.10453503 | 0.52672983 | 0.71336996 |
