## Supplementary material for "Alternative organelle targeting of OPA1 mediates fatty acid release from lipid droplets": Table S3

| Primers | Oligo sequence (5'-3') | Note |
|---|---|---|
| Opa1f-SalI | aaaa GTCGACatgtggcgactacgt | To amplify Opa1 from cDNA |
| Opa1r-BamHI | aaaa GGATCCCGtttctcctgatgaag | |
| 5'Opa1-158N/S | gaaaagcccttcctaGttcagaagaccttgt | Correct 158N to 158S |
| 3'Opa1-158N/S | acaaggtcttctgaaCtaggaagggcttttc | |
| 5'Opa1-S158N | gaaaagcccttcctaAttcagaagaccttgt | To generate Opa1 S158N |
| 3'Opa1-S158N | acaaggtcttctgaaTtaggaagggcttttc | |
| YFP Kozak remover f | GATCCACCGCTCGCCGTGAGCAAGGGCGAG | To remove the Kozak sequence in front of YFP in Opa1-YFP |
| YFP Kozak remover R | CTCGCCCTTGCTCACGGCGAGCGGTGGATC | |
| 5'Yfp-Opa1 ΔMTS | CGAATTCTGCAGTCGACatgttttggccagcaagattagct | To generate Opa1 ΔMTS-YFP |
| 3'Yfp-Opa1 ΔMTS | agctaatcttgctggccaaaacatGTCGACTGCAGAATTCG | |
| 5'Yfp-Opa1 iso2 | gatgagtatatcgattttggttctccggaagaaacg | To generate Opa1 isoform 2 |
| 3'Yfp-Opa1 iso2 | cgtttcttccggagaaccaaaatcgatatactcatc | |
| A195stop F | gaaacggcgtttagagcaTAacagatcgtggatctgaa | To generate Opa1 truncations ending at aa195 |
| A195stop R | ttcagatccacgatctgtTAtgctctaaacgccgtttc | |
| 5'Yfp-Opa1 127-c | CGAATTCTGCAGTCGACccggaccttagtgaatat | To generate Opa1 truncations starting at aa127 |
| 3'Yfp-Opa1 127-c | atattcactaaggtccggGTCGACTGCAGAATTCG | |
| 5'YFP-Opa1 157-c | CGAATTCTGCAGTCGACATGcctaGttcagaagacctt | To generate Opa1 truncations starting at aa157 |
| 3'YFP-Opa1 157-c | aaggtcttctgaaCtaggCATGTCGACTGCAGAATTCG | |
| gfp remover Tre lenti f | CCGGTCGCCACCATGGTGATGTctagactggacaag | To remove GFP from the 5' of YFP on Tre-iOpa1-YFP |
| gfp remover Tre lenti r | cttgtccagtctagACATCACCATGGTGGCGACCGG | |
| 5'lenti-iOpa1 | cgagcgtacgccaccatgtggcgactacgt | To generate TRE-OPA1-YFP-E3G |
| 3'Lenti-iOpa1 | TCTTTTATTCTAGCTaAGCTTACTTGTACAG | |
| 5' L156A | aaaattagaaaagccGCtcctaGttcagaagac | To introduce L156A mutation |
| 3' L156A | gtcttctgaaCtaggaGCggcttttctaatttt | |
| 5' F169A | aagttagcaccagacGCtgacaagattgttgaa | To introduce F169A mutation |
| 3' F169A | ttcaacaatcttgtcaGCgtctggtgctaactt | |
| 5' Opa1 G300E | ggagatcagagtgctgAaaagactagtgtgttg | To introduce G300E mutation |
| 3' Opa1 G300E | caacacactagtcttttTcagcactctgatctcc | |
| 5' Q297V | gttgtggttggagatGTGagtgctggaaagact | To introduce Q297V mutation |
| 3' Q297V | agtctttccagcactCACatctccaaccacaac | |
| ATGL f | ATTCCGTACGCCACCatgtttccccgcgag | To amplify ATGL from cDNA |
| ATGL R | aaaaACCGGTCCACCTCCcagccccagggccccgat | |
| ATGL S47A f | cacatctacggcgccgcggccggggcgctcacg | To generate ATGL dead mutant S47A |
| ATGL S47A r | cgtgagcgccccggccgcggcgccgtagatgtg | |
| CGI58F | aaaa AAGCTTaGCGGCGGAGGAGGAG | To amplify CGI58 from cDNA |
| CGI58R | aaaa GGATCCTCAGTCCACAGTGTCGCA | |
| exon4-gBlock | CTGCAGTCGACaccatggagaaaattagaaaagcccttcctaGttcagaagaccttgtaaagtta<br>gcaccagactttgacaagattgttgaaagccttagcttattgaaggactttttacctcaggtggcggaggg<br>ggatcAcGGGATCCACCG | synthetic fragment containing exon 4 |
| mhYFP-gBlock | CTGCAGTCGACaccatggagaaaattagaaaagcccttcctaGttcagaagaccttgtaaagtta<br>gcaccagactttgacaagattgttgaaagccttagcttattgaaggactttttacctcaggtggcggaggg<br>ggatcAcGGGATCCACCG | synthetic fragment containing mhYFP |
| OPA1 wMTS-gBlock | aaaaCTCGAGCTCAAGCTTCGAATTCTGCAGTCGACATGTGGGCTCTAG<br>CTGCTGCCGCTGTGGCCTGTGAGGTCTGCCAGTCTTTAGTGGCTCACA<br>GCTCTGGAATAGCTGGAAGTTTACCACTACAAGCTCTACATCTGGTTTC<br>ACGAAGCATTTATCATTCACATCATCCTACCTTAGCTCTTCAAGCTCCCC<br>AATTAAGGACATCCTTTCAGCAGTTCTCTTCTCTGACAAACCTTCCTTTA<br>GCTGCTCTGGCTTTCTCTCCAATTGCTTATGGCTACCAGCCTGCTGCTA<br>ATttttggccagcaagattagctacgagactcttaaaacttcgctatctcatactaggatcggctgttgggg<br>gtggctacacagccaaaaagacttttgatcagtggaaagatatgataccggaccttagtgaatataaatg<br>gattgtgcctgacattgtgtgggaaattgatgagtatatcgataaaa | synthetic fragment containing weakened OPA1 MTS |
| **RT-PCR primers** | | |
| hOpa1 RT-F | TACGCAAGATCATCTGCCAC | |
| hOpa1 RT-R | GAACGTGTCATCATCTCCCC | |
| hOpa1 RT 2-3F | GCTACACAGCCAAAAAGACT | |
| hOpa1 RT 3-5R | TCCGGAGAACCAAAATCGA | Combined with hOpa1 RT 2-3F to amplify fragments from isoform 2 (and exon 4 lacking isoforms) |
| hOpa1 RT 4-5R | CCGGAGAACCTGAGGTAAA | Combined with hOpa1 RT 2-3F to amplify fragment from isoform 1 (and exon 4 containing isoforms) |
| hGAPDH RT-F | GCCATCAATGACCCCTTCAT | |
| hGAPDH RT-R | GCTCCTGGAAGATGGTGATG | |
